# Social evolution and genetic interactions in the short and long term

**DOI:** 10.1101/010371

**Authors:** Jeremy Van Cleve

## Abstract

The evolution of social traits remains one of the most fascinating and feisty topics in evolutionary biology even after half a century of theoretical research. W. D. Hamilton shaped much of the field initially with his 1964 papers that laid out the foundation for understanding the effect of genetic relatedness on the evolution of social behavior. Early theoretical investigations revealed two critical assumptions required for Hamilton’s rule to hold in dynamical models: weak selection and additive genetic interactions. However, only recently have analytical approaches from population genetics and evolutionary game theory developed sufficiently so that social evolution can be studied under the joint action of selection, mutation, and genetic drift. We review how these approaches suggest two timescales for evolution under weak mutation: (i) a short-term timescale where evolution occurs between a finite set of alleles, and (ii) a long-term timescale where a continuum of alleles are possible and populations evolve continuously from one monomorphic trait to another. We show how Hamilton’s rule emerges from the short-term analysis under additivity and how non-additive genetic interactions can be accounted for more generally. This short-term approach reproduces, synthesizes, and generalizes many previous results including the one-third law from evolutionary game theory and risk dominance from economic game theory. Using the long-term approach, we illustrate how trait evolution can be described with a diffusion equation that is a stochastic analogue of the canonical equation of adaptive dynamics. Peaks in the stationary distribution of the diffusion capture classic notions of convergence stability from evolutionary game theory and generally depend on the additive genetic interactions inherent in Hamilton’s rule. Surprisingly, the peaks of the long-term stationary distribution can predict the effects of simple kinds of non-additive interactions. Additionally, the peaks capture both weak and strong effects of social payoffs in a manner difficult to replicate with the short-term approach. Together, the results from the short and long-term approaches suggest both how Hamilton’s insight may be robust in unexpected ways and how current analytical approaches can expand our understanding of social evolution far beyond Hamilton’s original work.

## 1. INTRODUCTION

The theory of evolution by natural selection as first fully elucidated by Darwin [27] is so profoundly elegant and comprehensive that truly new additions to theory have been extremely rare. In 1963, W. D. Hamilton began publishing his seminal work on how natural selection can shape social behavior [63–65], which is often either referred to as the theory of “kin selection” [106] or “inclusive fitness” [45]. It is a tribute to the importance of this work that upon his untimely death in 2000 Hamilton was called “one of the most influential Darwinian thinkers of our time” [39] and a candidate for the “most distinguished Darwinian since Darwin” [28].

In this article, we will review how the tools of population genetics and evolutionary game theory can be used to formalize Hamilton’s insight. We will begin with a summary of classic analyses of Hamilton’s approach and will then introduce the population genetic and game theoretic tools that currently provide a complete framework for studying social evolution under weak selection and weak mutation [98]. Using these tools, we will see how two general timescales for analysis emerge: a short-term timescale where evolution proceeds among a finite set of alleles, and a long-term timescale where populations evolve continuously among a continuum of alleles. These notions of short and long-term derive from a broader attempt to reconcile population genetic methods with evolutionary game theory [37, 68, 173].

Using the short-term approach, we show how genetic interactions between individuals [e.g. 127] can affect selection for cooperation in deme or group-structured populations [86]. These results extend previous analyses of stochastic evolution that have shown conditions such as “risk dominance” [14, 70, 78] and the “one-third law” [116, 120] to be important determinants of evolutionary stability. Using the substitution rate approach to long-term evolution [88, 165], we describe a diffusion equation that approximates the long-term change in monomorphic trait values. We show how peaks in the stationary distribution of this diffusion captures classic notions of evolutionary and convergence stability. Moreover, the location of these convergence stable states can be calculated using the classic direct-fitness approach of kin selection [132, 134, 155]. Applying this long-term approach to a simple non-additive social interaction, we find surprisingly that the long-term analysis can capture these non-additive effects even though the diffusion integrates over only additive interactions. Moreover, the long-term approach appears to reproduce results from some strong selection models, which suggests an unexpected robustness of the long-term diffusion. Together, the results from the short and long-term approaches reveal the usefulness of these approaches for integrating Hamilton’s original insight with recent results from population genetics and evolutionary game theory.

### 1.1. Hamilton’s rule

The core insight in Hamilton’s work is often summarized with his eponymous rule [64, 66]: an allele for a social behavior increases in frequency when the “inclusive fitness effect” is positive, namely

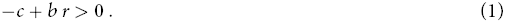

In Hamilton’s rule (1), *b* is the increase in fitness (benefit) of a social partner from the behavior of a focal individual, *c* is the decrease in fitness (cost) of a focal individual that performs the behavior, and *r* measures genetic relatedness between focal and recipient individuals [44]. More generally, − *c* is called the “direct fitness effect” and *b* the “indirect fitness effect”. Hamilton [64] initially emphasized that genetic relatedness is generated by a genealogical process that produces alleles identical by descent (IBD) among a group of socially interacting individuals. Another general definition of genetic relatedness says that it is the regression of the genotypes of social partners on the genotype of the focal individual [60, 66]. Hamilton’s rule crystalized the notion that natural selection depends both on the effect of an individual’s genes on its own fitness and also on the indirect effect of those genes on the fitness of social partners. Although Darwin [27], Fisher [42], and Haldane [62], among others, had expressed this idea in relation to the how evolution would lead one individual to sacrifice its fitness for another, Hamilton was the first to present a compelling framework applicable to social evolution more generally.

Within Hamilton’s inclusive fitness framework, behaviors that decrease the fitness of a focal individual (*c* > 0) but increase the fitness of social partners (*b* > 0) are “altruistic”. Well-known examples of altruism include worker sterility in eusocial insects [8], stalk cells that give up reproduction to disperse spore cells in *Dictyostelium discoideum* [145], and costly human warfare [67, 90]. Other behaviors can also be classified in Hamilton’s framework [64, and Table 2]: (i) behaviors are “mutualistic” when they increase the fitness of the focal individual and its social partners, (ii) “selfish” when they increase the fitness of the focal at the expense of the fitness of social partners, and (iii) “spiteful” when they decrease the fitness of both the focal individual and its social partners. Although there are other potential definitions of altruism and other behaviors [see 16, 82], Hamilton’s classification based on direct and indirect effects has proven useful for distinguishing different kinds of helping behaviors (mutualisms and altruisms) and for showing how different biological mechanisms can promote or inhibit the evolution of these behaviors [91, 174].

Though Hamilton’s approach was initially accepted among empiricists [181] and some theorists [106, 122], other theorists were concerned about the generality of the approach due to its emphasis on fitness maximization and optimality modeling [17, 79, 179]. Fitness maximization was viewed as untenable because examples where it is violated are well known [108]. Optimality models were additionally viewed with skepticism because, by neglecting gene frequency dynamics, they cannot study genetic polymorphisms; in effect, such models must assume that mutant alleles that invade a population also reach fixation. An initial wave of population genetic studies in response to these concerns showed that Hamilton’s rule was generally a correct mutant invasion condition so long as selection is weak and fitness interactions between individuals are additive [1, 17, 162, 163, 166]. However, these models were family structured where cooperation occurs between close relatives and could not address the applicability of Hamilton’s rule in populations with more generic structure, such as deme structure in island [182] and lattice models [84, 104, 105].

### 1.2. The Price equation and the individually-based approach

Part of the difficulty with the population genetic methods used to analyze family-structured models is that they use genotypes as state variables. This quickly increases the dimensionality of the model as the number of loci, family size, or demes increases and makes approximation difficult. An important alternative approach was introduced to population genetics by George Price with his eponymous equation [125, 126]. The Price equation uses the distribution of allele frequencies in each individual in the population as the set of state variables and tracks the first population-level moment of this distribution, which is the mean allele frequency. If **p**= (*p*_1_, ⋯, *p*_*N*_T__) represents the allele frequency distribution for *N*_T_ haploid individuals (*p*_*i*_ = 0 or 1 for individual *i*), the Price equation yields

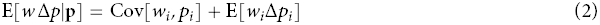

where E[*w* Δ*p*|**p**] is the expected change in mean allele frequency weighted by mean fitness *w* and conditional on **p** in the parental generation. The first term on the right hand side, the covariance between individual fitness *w*_*i*_ and allele frequency *p*_*i*_, measures the effect of selection on the change in mean allele frequency in the population. The second term, E[*w*_*i*_Δ*p*_*i*_], measures the effect of non-selective transmission forces, such as mutation and migration (and recombination for changes in genotype frequencies), on the change in mean allele frequency. When selection is the only force on allele frequencies and the population size remains fixed (*w* = 1), the Price equation simplifies to

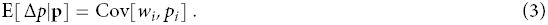

Calculating higher-order moments of the allele frequency distribution **p** is necessary to measure the exact dynamics of the distribution over time; thus, moment-based approaches like the Price equation are not necessarily more tractable than directly tracking genotype frequencies. However, an important observation about moment-based approaches is that they are readily amenable to approximation. When selection is weak relative to other forces such as recombination and migration, a kind of “separation of timescales” occurs where mean allele frequency dynamics converge very slowly and associations between alleles, such as linkage disequilibrium between loci [11, 85, 112] and *F*_ST_ between individuals in a deme [133, 137, 139, 169], converge much more quickly. Because of this separation of timescales, linkage disequilibrium, *F*_ST_, and other associations will converge to “quasi-equilibrium” (QE) values that are a function of mean allele frequencies. This means that the mean allele frequency dynamics can be expressed as a closed system of equations, which considerably simplifies analysis of multilocus systems or structured populations.

With respect to social evolution, the QE results for structured populations are particularly useful as they have helped to establish a rigorous basis for kin selection and Hamilton’s rule in populations with finite size, localized dispersal, or both [132, 134]. Initiated by the seminal work of François Rousset [131, 134], the mean allele frequency dynamics in this approach are calculated under weak selection and can be expressed as functions of *F*_ST_ and other between individual genetic associations evaluated under neutrality when selection is absent. In the simplest cases, this approach shows that Hamilton’s rule holds for weak selection and additive genetic interactions in populations with island-type structure [134] and family structure [138]. More generally, this approach produces analogues of Hamilton’s rule where the direction of selection is given by a sum of relatedness coefficients and fitness effects indexed by the spatial distance between a focal individual and its social partners [98, 132, 134] or by the demographic class (e.g., juvenile vs. adult or worker vs. queen in social insects) of the focal and its partners [135, 138]. It is this weak selection and QE approach that we will use to study genetic interactions and their affect on cooperation in deme-structured populations below.

### 1.3. Genetic drift, adaptive dynamics, and evolution in the short and long term

Another difficulty with the early analyses of kin selection and Hamilton’s rule in family structured populations was that those population genetic models could easily produce stable polymorphic equilibria [160, 162, 163], which made general predictions concerning the level of altruism or other social behaviors difficult. Generally, such equilibria are of intrinsic biological and mathematical interest since they illuminate stabilizing selection that can maintain genetic variation in levels of cooperation. In finite populations however, even alle-les under stabilizing selection either eventually go extinct or reach fixation due to genetic drift. If genetic drift is sufficiently strong relative to the rate of mutation *μ*, then the population will spend most of its time fixed for one of a set of possible alleles generated by mutation. This occurs for large *N*_T_ and small *μ* when

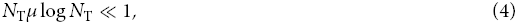

which can be arrived at heuristically by using the expected number of alleles in the population in an infinite alleles model under neutrality [80, 171, 172, 183]. Moreover, this condition ensures that any mutation that can arise will either fix or go extinct before another mutation arrives [18, 20]^1^; in other words, there are at most two alleles in the population at one time: a “resident” and a “mutant” that either goes extinct or fixes and becomes the new resident. This process of sequential substitution of alleles is called the “trait substitution sequence” [TSS; 19, 33] and is the fundamental biological model of adaptive dynamics [30, 53, 107] when population size goes to infinity, *N*_T_ → ∞. In addition, the TSS is often the implicit dynamical process behind many phenotypic models of kin selection and evolutionary game theory [29, 40, 98, 134, 155, 173, 175] that use an optimality criterion (i.e., fitness maximization) in search of an evolutionarily stable strategy (ESS).

On timescales short compared those required for generating phenotypic novelty, alleles generated by mutation in the TSS constitute a finite set, and the population “jumps” between alleles in this set as each allele invades and fixes. Assuming that it is possible to mutate from one allele to any other in the set through a sequence of zero or more intermediates (i.e., the mutation process is irreducible), the short-term process equilibrates to a stationary distribution λ among the fixation or monomorphic states. This short-term TSS corresponds to “short-term evolution” as defined by Eshel [36, 37] where a fixed set of genotypes are allowed to change frequency but new mutations outside this set do not occur. The length of time the population spends fixed for each allele is primarily determined by the likelihood each allele arises via mutation (*μ*) and fixes (fixation probability, *π*) in populations monomorphic for the other possible alleles [47]. Assuming weak selection, the QE approach discussed above can be used to calculate fixation probabilities even in spatially or demographically structured populations [132]. Together with the TSS condition (4), this allows a complete description of the stationary distribution of allelic states under the forces of selection, mutation, and genetic drift in the short-term.

For longer timescales, novel phenotypes are possible due to the invasion of mutations outside of a given finite set. For example, gene [102] and genome duplication [159], transposons [121], and lateral gene transfer [76] can generate novel physiological and ecological functions not possible with small changes in single genes. These processes suggest that the set of possible phenotypes may have a continuum of values over the long term. Suppose that for phenotype *z* the probability density of generating a mutant allele of phenotype *z* + *δ* is *u*(*δ*, *z*). If the support of *u*(*δ*, *z*) covers a fitness peak (i.e., it is possible to generate a mutant that resides exactly at the peak), then it is possible that the population will not only approach the peak, but it will spend most of its time fixed for a phenotype within a small neighborhood of the peak. This long-term TSS corresponds to the definition of “long-term evolution” by Eshel [36, 37] where invasion of new genotypes allow the population to approach phenotypic equilibria defined by the classic evolutionarily stable strategy (ESS) condition [38, 69, 101]. Without any assumptions on the distribution of mutational effects *u*(*δ*, *z*), the long-term TSS is described mathematically as a Markov jump process and is given by an integro-differential (master) equation [18–20, 88]. Often for the purpose of tractability, only small mutants are allowed in small intervals of time, which means that *u*(*δ*, *z*) is narrowly peaked around *z* and the population cannot make large jumps quickly. This assumption turns the jump process into a diffusion process [21] that is the stochastic analogue of the deterministic canonical equation of adaptive dynamics [19, 33]. The long-term TSS diffusion also has a stationary probability density, *ρ*(*z*), and the phenotypes located at peaks in that density correspond to equilibria obtained from classic ESS or adaptive dynamics analyses [88, 165]. Exactly as in the short-term TSS, weak selection can be used to calculate the fixation probabilities that determine *ρ*(*z*). This leads to a long-term stationary distribution of phenotypes that captures selection, mutation, and drift in spatially or demographically structured populations.

### 1.4. Putting it all together: Hamilton and social evolution in the short and long term

The two main assumptions above, weak selection and the TSS condition (4), allow us to describe the stationary density of transitions between a discrete set of phenotypes in the short term or among a continuum of phenotypes in the long-term. When there are only two types in the short term, cooperative and noncooperative, and the population is spatially structured or family structured, Hamilton’s rule in equation (1) is readily recovered when genetic interactions are additive [134, 138]. As we will see below, this is a result of comparing the stationary density of cooperative versus noncooperative types. Moreover, much of the recent work in evolutionary game theory that focuses on finite populations uses this same short-term TSS model to calculate a stationary distribution of types [e.g., 48, 72, 77, 116, 120, 141]. When there is a continuum of levels of cooperation in the long term, −*c* + *br* in Hamilton’s rule becomes the gradient of the potential function used to solve for the stationary density of the TSS diffusion [88]. Since −*c* + *br* can be thought to measure the change in inclusive fitness for additive genetic interactions [98, 134, 155], phenotypes at peaks in inclusive fitness (−*c* + *br* = 0) correspond to peaks in the stationary density. Thus, the long-term action of natural selection, *assuming additivity*, leads to a kind maximization of inclusive fitness, which supports the use of classic inclusive fitness analyses [however, see refs 97, 98, for difficulties in interpreting this result as broadly justifying “inclusive fitness maximization”].

### 1.5. Genetic interactions and non-additivity

If one is willing to assume weak selection, weak mutation relative to genetic drift (i.e., the TSS), and additive genetic interactions, then a direct application of Hamilton’s rule can be justified using the theoretical work discussed above. However, the ability to predict short and long term distributions of types under weak selection and the TSS is possible even when genetic interactions are non-additive. Non-additivity at the genetic level allows for interaction among alleles, within or between individuals. Within individuals, such interactions produce dominance and epistasis and between individuals they produce scenarios analogous to classic two-player games with pure strategies, such as the Hawk-Dove or Stag-Hunt games. Non-additive interactions are important because they produce frequency dependence in the sign of the change in allele frequency (eq. 3) even for weak selection [92, 119, 164]. In the case of social behavior, this implies that Hamilton’s rule becomes frequency dependent and no longer provides an unambiguous prediction of the effect of selection in either the short or the long term. Rather, applying the tools of QE and the TSS for non-additive interactions requires additional terms to account for higher-order genetic associations.

Once we calculate these additional terms, we determine the effect of non-additive interactions on the short-term stationary distribution of types in a given demographic context. Here, we apply the theory to Wright’s island model of population structure where there are *n* demes or groups each containing *N* haploid individuals [182] (*N*_T_ = *nN*). All groups are connected equally by migration at rate *m*. One of the important features of the Price equation approach is that it allows us to express the genetic associations (e.g., *F*_ST_) in terms of mean times to coalescence. Using results from coalescent theory to calculate the genetic associations, we replicate and generalize well known short-term results from inclusive fitness (the Taylor cancellation result [119, 151, 152]) and evolutionary game theory (the one-third law [116] and risk dominance [14, 70, 78]). Moreover, we show how changing the competitive environment (i.e., the degree of competition for local resources) changes these well known results, particularly in the presence of non-additive interactions.

The long-term approach, in contrast, only accounts for additive genetic interactions. Nevertheless, at least in social interactions with simple non-additive payoffs, the long-term approach remarkably reproduces results from the short-term approach that explicitly includes non-additive genetic interactions. We discuss a potential explanation for this power of the long-term approach, which suggests that three-way genetic interactions may be uniquely analytically tractable among possible non-additive interactions. Finally, we show how the long-term approach may capture some strong effects of social payoffs that the short-term approach neglects.

## 2. THEORY: SHORT-TERM EVOLUTION

### 2.1. Weak mutation,the TSS, and evolutionary success

Consider evolution in a population with total size, *N*_T_, where the population can be group structured (*n* groups of size *N*) or otherwise spatially structured with some pattern of migration between spatial locations. For the sake of simplicity, we assume that the population structure is homogenenous so that all individuals have equivalent effects on allele frequency change in the absence of natural selection [156]. Examples of homogeneous population structures include the island model and the stepping-stone and lattice models with a uniform migration rate. Inhomogeneous population structures include groups connected by dispersal via heterogeneous graphs, such as hub and spoke patterns, and require weighting different classes of individuals by their reproductive value [135, 147, 153].

Recall from section 1.3 that the short-term TSS requires considering only two alleles, which we label *A* and *a* where *p*_*i*_ measures the frequency of *A* in individual *i* (see Table 1 for a description of symbols used throughout this paper). Suppose that the mutation rate from *A* to *a* is *μ_a|A_* and *μ_A|a_* is the rate from *a* to *A*. We assume *μ* max *μ_A|a_ μ_a|A_* measures the overall strength of mutation. The weak mutation condition that defines the TSS, condition (4), is derived under the limit as *N*_T_ → ∞and →*μ* 0. In this limit, the TSS consists of the population jumping between states fixed for allele *A* and fixed for allele *a*. To represent the jump process between these two fixation states, we create a Markov chain for these two states with the following transition matrix

**Table 1:** Description of symbols.

| Symbol | Description |
| --- | --- |
| $N_T$ | Total population size |
| $N$ | Group (or deme) size |
| $n$ | Number of groups (or demes) |
| $m$ | Migration rate |
| $M$ | Population migration rate, $M = nNm/(n - 1)$ |
| $p_i$ ($p_{gi}$) | Frequency of allele $A$ in haploid individual $i$ (living in group $g$) |
| $p_g$ | Mean frequency of allele $A$ in group $g$ |
| $p(q)$ | Mean frequency of allele $A$ ($a$) in the population |
| $\mathbf{p} = (p_1, \dots, p_{N_T})$ | Vector of allele frequencies for the population |
| $\mu$ | Probability an offspring carries a mutant allele |
| $\mu_{A\vert a}$ ($\mu_{a\vert A}$) | Probability an offspring carries allele $a$ ($A$) with a parent carrying allele $A$ ($a$) |
| $z_i$ ($z_{gi}$) | Phenotype of individual $i$ (living in group $g$) |
| $\mathbf{z} = (z_1, \dots, z_{N_T})$ | Vector of phenotypes for the population |
| $\delta$ | Phenotypic deviation of mutant phenotype from resident phenotype $z$ |
| $u(\delta, z)$ | Distribution of mutant deviations $\delta$ (mutational effects) given a mutation from resident phenotype $z$ |
| $\sigma^2(z)$ | Second moment (raw variance) of the distribution of mutant deviations |
| $k(\delta, z)$ | Instantaneous rate of substitution of population monomorphic for trait $z$ with one monomorphic for trait $z + \delta$ |
| $\rho(z, t)$ ($\rho(z)$) | Probability density that population is monomorphic for trait $z$ at time $t$ ($t \rightarrow \infty$) |
| $w_i$ ($w_{gi}$) | Fitness of individual $i$ (living in group $g$) |
| $w$ | Mean fitness in the population |
| $b$ | Fitness benefit to the focal individual of the expression of allele $A$ in a social partner |
| $c$ | Fitness cost of the expression of allele $A$ in the focal individual |
| $f_{ig}, f_g, f$ | Fertility of individual $i$ in group $g$, mean fertility in group $g$, mean fertility in the population |
| $\pi_{A\vert a}$ ($\pi_{a\vert A}$) | Probability of a single mutant $A$ ($a$) allele fixing in a population of $N_T - 1$ $a$ ($A$) alleles |
| $\pi^\circ$ ($\pi^\circ(z)$) | Probability of a neutral allele (in a population resident for phenotype $z$) fixing in the population |
| $S(z)$ | Derivative of fixation probability with respect to $\delta$ and evaluated at $\delta = 0$; also called the “selection gradient” |
| $\omega$ | Selection strength |
| $s_{i,k_1 \dots k_d}$ | Selection coefficient for individual $i$ of the frequency of $A$ in individuals $k_1, \dots, k_d$ |
| $P_i(\mathbf{p})$ | Sum of selection coefficients for individual $i$ times their respective allele frequency products |
| $\mathbf{P}(\mathbf{p}) = (P_1(\mathbf{p}), \dots, P_{N_T}(\mathbf{p}))$ | Vector of sums of selection coefficients and allele frequency products |
| $E^\circ[T_{ik_1 \dots k_d}]$ | Expected coalescence time of alleles in individuals $i$ and $k_1$ through $k_d$ under neutrality |
| $Q_{ij}$ | Probability of identity by descent between alleles in individual $i$ and $j$ |
| $r$ | Genetic relatedness |
| $\kappa$ | Scaled relatedness coefficient |
| $k$ | Selection gradient proportionality constant |

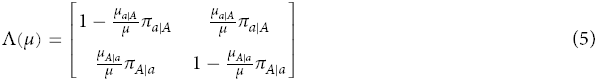

where *π_A|a_* and *π_a|A_* are the probabilities that alleles *A* and *a*, respectively, reach fixation starting from an initial frequency of 1 *N*_T_ in a population where the other allele has frequency 1–1 *N*_T_. Rescaling the mutation rates by the overall rate *μ* allows a nontrivial stationary distribution of the Markov chain (i.e., the left eigenvector of Λ(*μ*)) as *μ* → 0. Inspired by ideas in large deviations theory [46], Fudenberg and Imhof [47] show that the stationary distribution λ of the TSS as *μ* → 0 is simply the stationary distribution of the Markov chain in (5) in the limit as *μ* → 0. Thus, instead of having to calculate the stationary distribution of the complex stochastic process with many different possible population states, we need only calculate the stationary distribution of the much simpler “embedded” chain composed of fixation states. Calculating the stationary distribution using the embedded chain yields

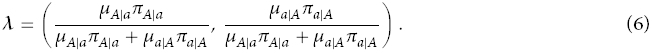

If we are interested only in the effect of selection on the stationary distribution, we can assume that the mutation rates are symmetric, *μ_A|a_* = *μ_a|_*= *μ*. In this case, the expected frequency of allele *A* in the population at stationarity, which we write as E[*p*], becomes in the limit as the mutation rate goes to zero

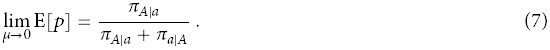

An intuitive condition for the evolutionary success of allele *A* relative to allele *a* is that *A* is more common at stationarity, or

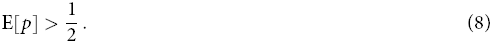

Using equation (7), condition (8) is equivalent to

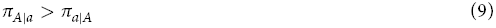

when *μ* → 0, which means we need only compare complementary fixation probabilities in order to determine which allele is “favored” by natural selection [7, 48]. This condition on fixation probabilities is the evolutionary success condition that we will use to derive Hamilton’s rule (eq. 1).

### 2.2. Fixation probability and the Price equation

Calculating the fixation probabilities in a model with arbitrarily complex demography or spatial structure can be daunting if not impossible. Thus, our next aim is to show how to connect fixation probabilities to the Price equation, which will make it straightforward to use weak selection and QE results. Suppose that *p*(*t*) is the mean frequency of allele *A* at time *t*. Following recent methods [95, 100, 131], we can write the fixation probability as

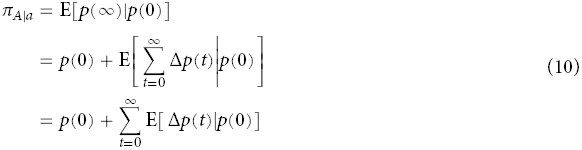

where Δ*p*(*t*) = *p*(*t* + 1) - *p*(*t*) and we can exchange the expectation and the infinite sum in the last line because the Markov chain converges in mean [100]. Expanding the sum in (10) by conditioning on all possible population states **p**(*t*) yields

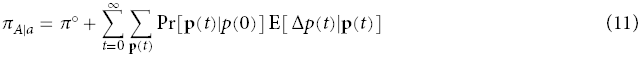

where we have used the fact that the fixation probability of a neutral allele, *π*°, is its initial frequency *p*(0). The second term in the sum, E[ Δ*p*(*t*)|**p**(*t*)], is exactly the left-hand side of the Price equation (3).

### 2.3. Fixation probability under weak selection

The most straightforward way to calculate the probability of fixation *π_A|a_* assuming weak selection is to Taylor expand *π_A|a_* in terms of a parameter that measures the strength of selection [see: 95, 100, 131], which we call *ω*. This expansion is simply

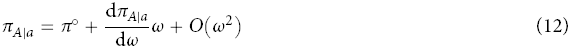

where the derivative 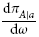 is evaluated under neutrality (*ω* = 0). Using equation (11), we can calculate the derivative of the fixation probability under neutrality as

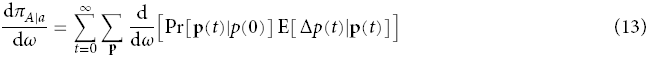

where the exchange of derivative and the limit is justified provided the derivatives converge uniformly [see Appendix of 100, for such a proof]. Expanding the derivative in the sum in (13) using the chain rule yields

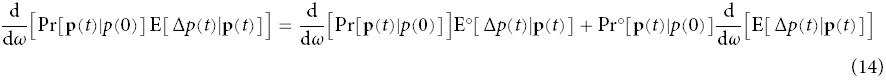

with the symbol ° indicating evaluation of an expectation or probability in the neutral case when *ω* = 0. The first term on the right hand side of (14) is zero since the expected change in allele frequency under neutrality is zero for homogeneous population structures. Simplifying equation (13) with this fact yields

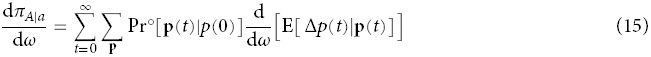

which we can write as

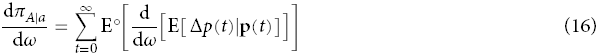

where E° implies expectation over the neutral realizations of **p** given an initial frequency of *A* of *p*(0).

In order to evaluate the derivative of the fixation probability in equation (16), we need the derivative of the expected change in mean allele frequency. This is a quantity that is relatively simple to calculate since all one needs is the first-order term in an expansion of E[ Δ*p*(*t*)|**p**(*t*)] in terms of selection strength *ω*. To obtain this expansion, we first expand the fitness of the focal individual *i* in terms of selection strength. Without loss of generality, the fitness of individual *i* is

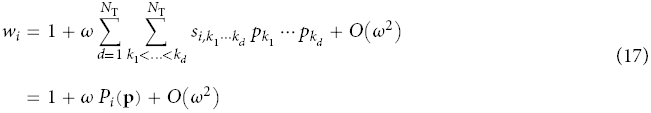

where *s*_*i*,1_, ⋯, *s*_*i*,1⋯*N*_T__ are often called “selection coefficients” [11, 85] and are time independent and *P*_*i*_(**p**) is the polynomial given by the summations in the first line. The *d* = 1 term in the summation yields the “additive” fitness components where selection coefficients *s*_*i,j*_ are multiplied *p*_*j*_. Terms in the summation *d* > 1 are “non-additive” fitness components since those selection coefficients are multiplied by products of allele frequencies. Given that mean fitness is equal to one (which is true for populations of fixed size *N*_T_), *P*_*i*_(**p**) must satisfy to first order in *ω*

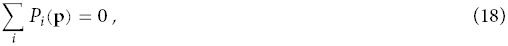

which also implies that the sum of the selection coefficients must be zero by setting **p** = **1** in (18); this constraint on the sum of the selection coefficients is a common feature of weak selection models with a fixed demography [e.g.: 95, 134].

Using the expression for fitness in (17), the change in mean allele frequency from the Price equation (3) becomes

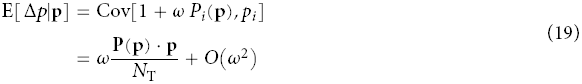

where **P**(**p**) = (*P*_1_(**p**), ⋯, *P*_*N*_T__ (**p**)) and **P**(**p**) · **p** is the scalar product of the two vectors. For example, if in a single panmictic population allele *A* confers a fitness advantage of *ω* for every individual that has it, then *w*_*i*_ = 1 + *ω*(*p*_*i*_ – *p*) and

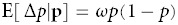

which is the standard result for an advantageous allele under weak selection [41]. Taking the first-order term from (19) and inserting it in equation (16) produces

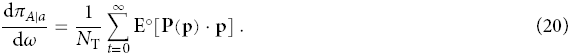

Expanding (20) into the first-order Taylor series for the probability of fixation in (12) yields

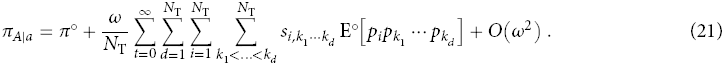

At this point, even though we have not explicitly specified the effect of gene expression on fitness or the population structure, we can see how additive and non-additive effects of selection affect fixation probability. The additive terms, *d* = 1 in the above sum, depend on expected pairs of allele frequency, E[*p*_*i*_*p*_*j*_]. These expected pairs are essentially probabilities of genetic identity between two different individuals in the population. Thus, measures of average pairwise genetic identity within a structured population, such as Wright’s *F*_ST_, are natural statistics to use when considering the effect of selection due to additive genetic interactions. The non-additive terms, *d* > 1, contribute expected (*d* + 1)-order products of allele frequency and thus require higher order statistics than *F*_ST_.

In order to better interpret the expression for fixation probability in (21), which contains a difficult infinite sum over time, we follow the argument given in Rousset [131] and expanded in Lessard and Ladret [100] and Lehmann and Rousset [95] that interprets the expected allele frequency products in terms of coalescence probabilities. Recall from the TSS that in a population composed of allele *a*, a single *A* mutation will arise and either fix or go extinct. In this case, the expected allele frequency product, E°[*p*_*i*_*p*_*k*_1__ ⋯ *p*_*k*_*d*__], is the probability that individuals *i* and *k*_1_ through *k*_*d*_ all have allele *A* at some future time *t*. Going backwards in time, this probability is equivalent to the probability that those lineages coalesce before time *t*, Pr°[*T*_*ik*_1_⋯*k*_*d*__ ≤ *t*], times the probability that the ancestral lineage is allele *A*, which is the initial frequency *p*(0) = *π*° = 1/*N*_T_. Writing Pr°[*T*_*ik*_1_⋯*k*_*d*__ ≤ *t*] as 1 − Pr°[*T*_*ik*_1_⋯*k*_*d*__ > *t*] and using the fact that the selection coefficients sum to zero (eq. 18), equation (21) becomes

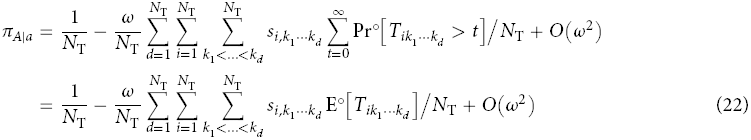

which matches previous results [eq. 15 in 131; eqs. 59 and 61 in 100]. Equation (22) says that the effect of selection on the fixation probability of allele *A* is simply a sum of selection coefficients and expected coalescence times under neutrality, E°[*T*_*ik*_1_⋯*k*_*d*__]. One advantage of expressing the fixation probability in terms of coalescence times is that results from coalescence theory [170] can be used, which is the approach we utilize when we apply these methods to a population with island-type structure.

The condition for allele *A* to be more common at stationarity than allele *a*, condition (9), requires both fixation probabilities *π_A|a_* and *π_a|A_*. Expanding *π_a|A_* under weak selection can be accomplished using the same reasoning above for equation (22). First, observe that that the expected change in the frequency of the *a* allele is E[Δ*q*(*t*)] = − E[Δ*p*(*t*)] where *q* = 1 − *p*. This implies that 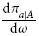 is simply the negative of the first-order term in equation (21) *except* that E°[*p*_*i*_*p*_*k*_1__ ⋯ *p*_*k*_*d*__] is evaluated under a neutral process where the initial frequency of *a* is 1/*N*_T_ (rather than 1 - 1/*N*_T_ as was the case for *π_A|a_*). In calculating *π_A|a_*, E°[*p*_*i*_*p*_*k*_1__ ⋯ *p*_*k*_*d*__] was interpreted as a coalescence probability times the initial frequency of *A*; analogously, we can replace each *p*_*i*_ with 1 − *q*_*i*_ and interpret the products *q*_*i*_*q*_*k*_1__ ⋯ *q*_*k*_*d*__ in E°[(1 − *q*_*i*_)(1 − *q*_*k*_1__) ⋯ (1 − *q*_*k*_*d*__)] as a coalescence probabilities, Pr°[*T*_*ik*_1_⋯*k*_*d*__ ≤ *t*], times the initial frequency of *a*. Calculating *π_a|A_* using the analogue of (21), expanding the expected products of *q*_*i*_, and summing over time yields

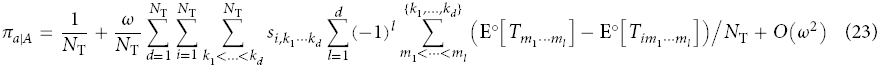

where the last sum on the right-hand side is over all *m*_1_ < ⋯ < *m*_*l*_ drawn from the set {*k*_1_, ⋯, *k*_*d*_} and E°[*T*_*k*_*j*__] = 0 for any single lineage *k*_*j*_ by definition.

With expressions from both fixation probabilities, *π_A|a_* from equation (22) and *π_a|A_* from equation (23), we can combine them to complete the condition for allele *A* to be more common than *a* at stationarity, *π_A|a_ > π_a|A_*, which becomes

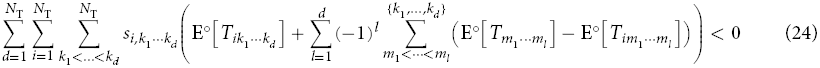

to first order in *ω*. This is the main condition for evolutionary success for the short-term TSS. Given a population where the demography and the relationship between gene expression and fitness are known (i.e., the selection coefficients *s*_*i,k*_1_⋯*k*_*d*__ are known), all that remains is to calculate expected coalescence times under neutrality, which will be a function of the demography. For models of cooperative behavior, condition (24) generates both Hamilton’s rule when genetic interactions are additive (*d* = 1) and the risk-dominance condition when interactions are non-additive (*d* > 1).

### 2.4. Additive genetic interactions and evolutionary success

The effect of selection on fixation probability given in (22) can be arbitrary complex if individual fitness depends on any combination of genotypes (i.e., products of allele frequency) of individuals in the population. In the simplest social interactions, fitness only depends additively on the genotype of focal individual and the genotypes of other individuals in the population, which means *d* = 1 in equation (17). Using *d* = 1 in the expression for *π_A|a_* in (22), we find that

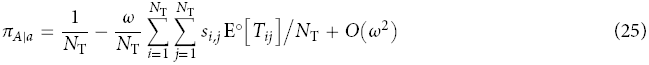

Similarly, applying *d* = 1 for the expression for *π_a|A_* produces

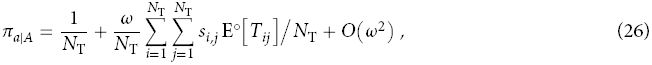

which is identical to equation (25) except the first order term has the opposite sign. Adding equations (25) and (26) immediately yields

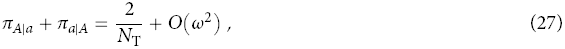

which transforms the evolutionary success condition *π_A|a_* > *π_a|A_* to (after neglecting *O*(*ω*^2^) terms)

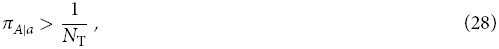

which is the standard condition for type *A* to be advantageous compared to a neutral allele. Inserting the first-order expansion for *π_A|a_* in equation (12) in condition (28) produces another evolutionary success condition for allele *A*:

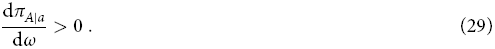

Thus, additive genetic interactions simplify the analysis of evolutionary success considerably since we have to evaluate only one fixation probability, *π_A|a_*, or its derivative with respect to the strength of selection. Conversely, analyses that only use condition (28) and compare the fixation probability of one type versus fixation under neutrality are correct predictors evolutionary success *only* when interactions are additive. For example, the one-third law from evolutionary game theory is an evaluation of condition (28) in a finite population where individuals interact in a pairwise manner and the payoffs from their interactions can produce non-additive genetic interactions [116, 120]. In effect, the one-third law then yields a measure of the relative stability of a population fixed for allele *a*, but does not provide information about the stability of the population when fixed for allele *A* [see Fig. 2b in 116] unless genetic interactions are additive.

### 2.5. Weak effect mutations and continuous phenotypes

An important case where genetic interactions are additive occurs when the difference between the phenotypes produced by alleles *A* and *a* is small and phenotypes are allowed to take a continuum of values. Suppose that the phenotype of type *a* is *z* and that of type *A* is *z* + *δ* where *δ* is called the phenotypic deviation. Further, let the phenotype of individual *i* be *z*_*i*_ = *z* + *δp_i_*, and the vector **z** = **z**(**p**) = (*z*_1_, ⋯, *z*_*N*_T__) contain the phenotypes for the whole population. Since phenotype is a continuous variable, we assume that the fitness of each individual *i*, *w*_*i*_(**z**), is a differentiable function of phenotype [p. 41 in ref 26]. This also implies, using equations (3) and (11), that the fixation probability a single mutant with trait *z* + *δ* in a population with resident trait *z*, denoted *π*(*δ*, *z*), is differentiable. When the phenotypic deviation *δ* is small (weak effect mutations), we can ignore terms *O*(*δ*^2^) and individual fitness can be written as a Taylor series in *δ*:

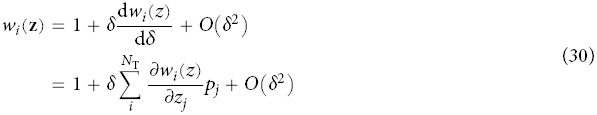

where 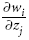 are evaluated at *δ* = 0 (written as 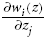)when the population is fixed for allele *a* and *w*_*i*_(*z*) = 1. Comparing equation (30) to the expression for fitness when phenotypes are discrete (eq. 17) reveals that the phenotypic deviation *δ* is analogous to the selection strength *ω* and that the derivatives of fitness with respect to phenotype, 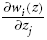 are equivalent to additive selection coefficients. Thus, so-called “*δ*-weak” selection [176] implies additivity of fitness effects and allele frequency, which is well known in the literature [e.g.: 132, 150].

Applying the fitness function in (30) to the fixation probability equation (25) produces

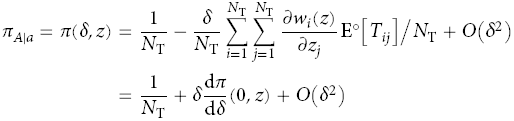

where

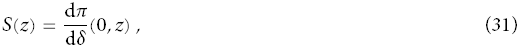

is often called the the phenotypic “selection gradient” in adaptive dynamics [53, 99], inclusive fitness theory [96] and quantitative genetics [87]. The “gradient” terminology derives from the fact that *S*(*z*) measures the direction of selection on the phenotype with respect to fixation probability: a positive (negative) selection gradient implies that mutants with positive (negative) *δ* will have a higher (lower) chance of fixing than the resident type. The zeros of the selection gradient, which correspond to extrema of the fixation probability, are candidate evolutionary equilibria. We will show this to be the case though once we have described evolution under the long-term TSS in section 4.

### 2.6. Coalescence time and identity by descent

So far, we have shown how the fixation probability of an allele with effects on social behavior depends on mean coalescence times (eq. 22). However, the relatedness term in Hamilton’s rule is often expressed as a function of probabilities of identity by descent [64, 66]. Translating between mean coalescence times and probabilities of identity by descent is possible using an argument first presented by Slatkin [143]. Suppose that *Q*_*ij*_ represents the expected probability of identity by descent between alleles *i* and *j*. Since mutations are distributed independently on a neutral genealogy [170] and IBD requires that the alleles not mutate before coalescing, the IBD probability can be expressed as

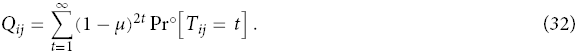

Since the TSS assumes weak mutation, we can ignore terms *O*(*μ*^2^) and rewrite equation (32) as

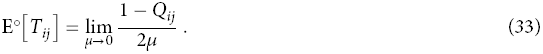

We use the limit as *μ* → 0 in equation (33) since we derived the fixation probabilities and their dependence on coalescence time under the TSS assumption that new mutation is not possible until the old mutation either fixes or goes extinct. Equation (33) only pertains to pairwise coalescence times and IBD probabilities, so we can only apply it to additive genetic interactions. The relationship between three-way (and more generally *d*-way) coalescence times and IBD probabilities is more complex and deserves further study.

Applying the relationship between pairwise coalescence times and IBD probabilities in (33) to the selection gradient in (31) in the case of *δ*-weak selection yields

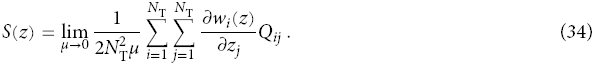

This expression was first obtained by Rousset and Billiard [134], and an analogous derivation was presented by Rousset [131]. For models with simple population structure (homogenous structures [119, 156] like the island [182] or stepping-stone [84] models) and simple demography, the IBD probabilities *Q*_*ij*_ are relatively easy to obtain in the low mutation limit. The fitness function *w*_*i*_(**z**) depends on nature of the social interactions as well as on the demography and population structure.

### 2.7. Inclusive fitness effect and Hamilton’s rule

Essentially, the right hand side of (34) is a measure of how expression of allele *A* affects inclusive fitness [132, 134]; fitness effects are given by the derivative of the fitness of individual *i* with respective to the phenotype of individual *j*, and each effect is weighted by the likelihood that individuals *i* and *j* share alleles IBD. Applying the evolutionary success condition *S*(*z*) > 0 to (34) yields a Hamilton-type rule,

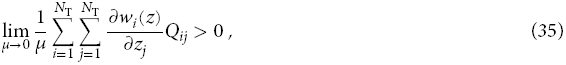

where the population structure and fitness functions haven’t been specified. In order to obtain the classic form of Hamilton’s rule from condition (1), we make some simplifying assumptions about the social interaction and population structure.

Suppose that there is a homogeneous population structure with *n* groups each containing *N* haploid individuals (*N*_T_ = *nN*) and equal migration between all groups (i.e., an island model). This implies that we need only to track two kinds of IBD probabilities, *Q*_0_, which measures the chance that two alleles drawn from different individuals in the same group are IBD, and *Q*_1_, which is the probability that two alleles from different groups are IBD. Individuals socially interact within their group, but social effects between groups also occur due to differential productivity of groups (i.e., “hard selection” [23]). Since the fitness derivatives 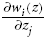 are evaluated at *Δ* = 0 where all individual behave the same (as if they express the *a* allele), there are only three different fitness derivatives: (i) individuals *i* and *j* are the same individual, and 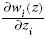 is the effect of the individual’s behavior on itself, which we call the “cost” or −*c*; (ii) individuals *i* and *j* live in the same group, and 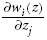 is the effect on the individual due to its group mate’s behavior, which we call the “benefit” or *b*/(*N*− 1) (for each of the *N* − 1 group mates); and (iii) individuals *i* and *j* live in different groups where we can set 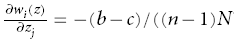 since the selection coefficients must sum to zero (eq. 18). Putting these expression into equation (35) and simplifying produces

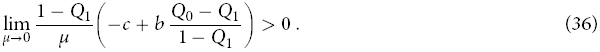

Typically, in finite populations (*N*_T_ < ∞), the IBD probabilities *Q*_0_ and *Q*_1_ will go to one as the mutation rate goes to zero since one lineage will eventually fix in the population. In these cases, the IBD probabilities can often be expressed as 1 − *μ O*(1) [132], which suggests that the first term in (36) has a positive limit as *μ* → 0. The ratio multiplying the benefit *b* turns out to be Wright’s *F*_ST_, which will also have a positive limit under low mutation. Setting the relatedness to 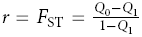 and simplifying, we then obtain Hamilton’s rule for this population

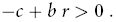

where the left-hand side of the inequality is the inclusive fitness effect. Using Slatkin’s formula in (33), we can equivalently write relatedness in this context as [143]

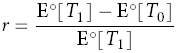

where E°[*T*_0_] and E°[*T*_1_] are the expected coalescence times between alleles sampled in the same and different groups, respectively.

In his derivation of the inclusive fitness effect, Hamilton [66] began by directly applying the Price equation to the change in the frequency of an allele affecting social behavior. His relatedness coefficient *r* in that context is a regression coefficient for the frequency of the allele in a focal individual as a function of the frequency of the allele in social partners. This is equivalent to assuming that E[*p*_*j*_|*p*_*i*_] = *r p*_*i*_ + (1 − *r*)*p* for focal individual *i*, social partner *j*, and population-wide frequency *p* [44, 66, 132, 134]. In an island model like the one used in equation (36), this means that a social partner in the group either shares ancestry with the focal individual in proportion to *r* and has the same allele frequency as the focal or doesn’t share ancestry and has the population-wide frequency. Relatedness using this regression definition is

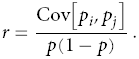

The covariance term above can be written as Cov[*p*_*i*_, *p*_*j*_] = E[*p*_*i*_*p*_*j*_] - *p*2 where E[*p*_*i*_*p*_*j*_] is the probability that both individuals *i* and *j* in the same group have allele *A*. Since E[*p*_*i*_*p*_*j*_] = *Q*_0_*p* + (1 − *Q*_*o*_)*p*2, the regression defintion of relatedness becomes *r* = *Q*_0_ in this example. This matches 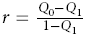 from equation (36) since the regression assumption implies that individuals in different demes (individuals outside the social interaction) share no ancestry, which means *Q*_1_ = 0. One way this can occur is if the number of groups is infinite (*n* → ∞). There are two important points about Hamilton’s rule that are illuminated by the derivation in equation (36). First, even though relatedness can be defined as a regression coefficient and thus might appear to capture only a statistical snapshot of population structure, the definition above in terms of coalescence times or IBD probabilities emphasizes that relatedness is a multigenerational concept. Genetic associations between alleles are created by the long term effects of mating, reproduction, dispersal, and survival on genealogy, which can be measured with coalescence times. These associations, through coalescence times or IBD probabilities, will generally depend explicitly on the demography and population structure. More broadly, this also implies that the inclusive fitness effect in Hamilton’s rule captures multigenerational effects of selection on allele frequency change.

The second point is that the cost and benefit terms are effects on *fitness* as measured over the whole lifecycle. This implies that fitness effects will be functions of demographic parameters (such as population size and migration rate) and of fertility or survival payoffs from the social interaction. In order to determine the signs of −*c* and *b*, which are required for classifying a behavior as altruistic or not (see Table 2), the demography and the effect of social behavior on both fertility and survival must be specified.

**Table 2:** Definitions of social behavior using Hamilton’s rule (eq. 1)

| Effect on<br>focal ($-c$) | Effect on social partner ($b$) | |
| --- | --- | --- |
|  | + | - |
| + | Mutualism | Selfishness |
| - | Altruism | Spite |

## 3. APPLICATION: SOCIAL GAMES IN ISLAND-TYPE POPULATIONS

To recap a bit, we have shown above how weak mutation and the TSS allow a simple criterion for evolutionary success, *π_A|a_* > *π_a|A_*, in the short term. When selection is weak and genetic interactions are additive, this simplifies the condition for evolutionary success considerably and we recover a measure of inclusive fitness in (34). When non-additive genetic interactions are important and selection is still weak, the evolutionary success condition is given more generally by condition (24).

There are many biological scenarios where non-additive interactions are important, and the simplest one that is often invoked is a two-player game. Each individual plays one of two pure strategies, “cooperation” or “noncooperation”, with a social partner where the payoffs for the game are given in Table 3. When two individuals are noncooperators, they receive no payoff. If one does not cooperate and the other cooperates, the cooperator receives −*C* and the noncooperator receives *B*. When two individuals cooperate, they each receive payoff *B* - *C* + *D* where *D* is a measure of non-additivity or “synergy”. The strategy names and payoffs are inspired by the Prisoner’s Dilemma game [12] where *B* and *C* are positive and *D* ≥ 0, though we will allow the parameters to take negative values as well in order to study other games like the Stag-Hunt [142]. In the simplest case, the strategies are fixed by the genotype of the individual where individuals bearing the *A* allele always cooperate and individuals bearing *a* always do not cooperate. This means there is no phenotypic plasticity that might result from changing strategies over repeated interactions (e.g., reciprocity [9, 161] or responsiveness [2, 3]).

**Table 3:** Payoffs for the cooperation (coop) allele, *A*, and noncooperation (noncoop) allele, *a*, in the social game.

| Focal<br>individual | Social partner |  |
| --- | --- | --- |
| | $A \mid \text{coop}$ | $a \mid \text{noncoop}$ |
| $A \mid \text{coop}$ | $B - C + D$ | $-C$ |
| $a \mid \text{noncoop}$ | $B$ | $0$ |

We assume an island-type population structure where *n* groups of *N* haploids (*N*_T_ = *nN*) are connected each by migration rate *m*. Generations are non-overlapping. The frequency of the cooperation allele *A* in individual *i* in group *g* is *p*_*gi*_, the mean frequency in group *g* is *p*_*g*_ including individual *i* and *p*_*g\i*_ excluding *i*, and *p* is the mean frequency in the whole population. Individuals choose social partners at random and the mean payoff from the social interactions in the group determines how each individual’s fertility differs from the baseline of one. Given these assumptions, the fertility of individual *i* in group *g* is 1 + *ωf_gi_* where

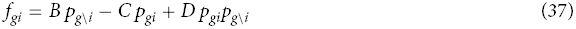

and *B* and *C* are additive and *D* non-additive in allele frequency. This fertility function represents not only the two-player case but also the *n*-player case within the group when payoff is an additive function of the frequency of other player types in the group [165, Appendix C].

### 3.1. Baseline demography: hard selection

We begin with a model of hard selection (see section 2.7) with groups producing different numbers of migrants depending on their composition. The fitness of individual *i* in group *g* can be written as

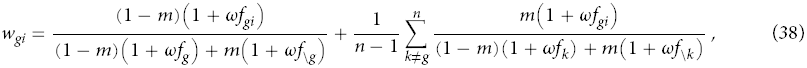

where 1 + *ωf*_\*g*_ is the mean fertility in the population excluding group *g*, or to first order in *ω* as

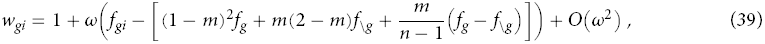

which takes the same form as equation (17). All that remains in order to calculate the fixation probabilities in equations (22) and (23) are the expected coalescence times under neutrality.

Fortunately, coalescence times in structure populations are a well-studied topic [73, 115, 168, 178] in coalescent theory where the process often considered is called the structured coalescent [113, 114]. As applied to the island model, the structured coalescent usually assumes that the migration rate *m* is *O*(1/*N*) and *nNm*/(*n* − 1) → *M* as *N* → ∞. This impies that during a small interval of time, either two lineages within a group can coalesce or one lineage can migrate from one group to another, but more than one such event does not occur. For two lineages, either both can be in the same group with configuration (0, 1) or each can be in different groups with configuration (2, 0) = (2) where the first element of the configuration is the number of groups with a single lineage, the second element is the number of groups with two lineages, etc (e.g., see [168]). We denote the expected coalescence times of these configurations by E°[*T*_(0,1)_] and E°[*T*_(2)_], respectively. With synergistic payoffs (*D* ≠ 0) that create non-additive fitness effects, we also have to track three lineage samples. For three lineages, there are three possible configurations: (3), (1, 1), and (0, 0, 1). Using the master equation for the continuous-time Markov process that describes coalescence (eq. 2.8 in [115]), a system of equations for the five expected coalescence times can be constructed (Wakeley [168] describes the method nicely); these equations are given in section 5.1 of [86], and we only provide the solutions here (time in absolute units):

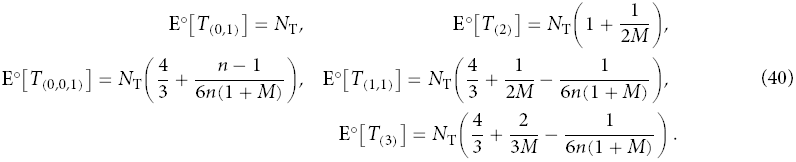

Note that evaluating a more complex fitness function with higher-order frequency dependence would require calculating expected coalescence times for four lineage or larger samples. The number of configurations and equations grows quickly as the lineage sample size increases, which makes this method cumbersome for complex fitness functions. Working on the related problem of calculating the total length of the coalescent genealogy, Wakeley [168] provided a method to calculate expected coalescence times for arbitrarily large samples so long as the number of groups *n* is large; this suggests that an analogous method might work for coalescence times that could be used to calculate fixation probabilities.

Applying the fitness function from (39) and the coalescence times in (40) to equation (22), we calculate the probability allele *A* (the cooperation allele) fixes in a population of *a* (the noncooperation allele) as

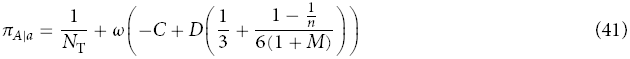

and the fixation probability of *a* in a population of *A* as

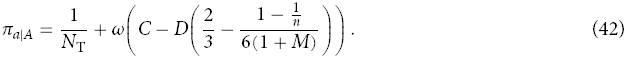

First derived by Ladret and Lessard [86, eq. 29], these expressions are correct to first order in selection strength *ω* and zeroth order in *O*(1/*N*) since our coalescence times assume that *N* → ∞. These two fixation probabilities are the first main result of this section and produce a few important observations. First, they reproduce the classic cancellation result from Taylor [151, 152, 156] for additive genetic interactions (*D* = 0). Taylor’s result says that the benefits of cooperation are exactly balanced out by the effect of competition between related individuals within a group (so-called “kin competition” or “local competition”) when population structure is “homogenous” [156] and generations do not overlap. In our model notation, the cancellation result implies that the benefit to others *B* will cancel out of the fixation probability expressions; equations (41) and (42) show that this indeed does occur. Thus, when interactions are additive (*D* = 0), the cooperation allele *A* fixes with a probability greater than the neutral probability *π*° = 1/*N*_T_ (eq. 28) only when there is negative direct cost (i.e., cooperation is directly beneficial). Likewise, *π_a|A_* in equation (42) shows that the noncooperation allele *a* is advantageous when the cost is positive. In an extension of the cancellation result, Ohtsuki [119] shows (for *n* → ∞) that positive that synergy that does not change the structure of the game, 0 < *D* < *C*, cannot not result in positive selection for cooperation. Equations (41) and (42) reproduce this result since even the strongest population structure, *M* → 0, results in cooperation fixing more likely than chance only when *D* > 2*C*.

The expression for *π_A|a_* in equation (41) also produces the one-third law from evolutionary game theory [116]. The one-third law says that as *N* → ∞ in a single panmictic population, the cooperation allele *A* fixes with a probability greater than chance when the mixed strategy equilibrium of the game in Table 3 is less than one third. If an opponent cooperates with probability *z* and does not cooperate with probability 1 − *z*, the mixed strategy equilibrium is the value of *z* where an individual does equally well against the opponent by either cooperating or not cooperating [75]. Using the payoffs from Table 3, the mixed strategy equilibrium equation is *z*\*(*B* − *C* + *D*) − (1 − *z*\*)*C* = *z*\**B*, which yields *z*\* = *C*/*D*. Thus, the one-third law translates to

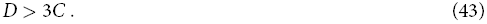

We can immediately recover the one-third law from *π_A|a_* > 1/*N*_T_ by taking the high migration limit *M* → ∞ in (41), which results in an unstructured population. The complementary condition for fixation of the noncooperation allele *a*, *π_a|A_* > 1/*N*_T_, becomes *D* < 3*C*/2 in the high migration limit. In contrast, population structure is at its strongest in the low migration limit when *M* → 0 and when the number of groups is large, *n* → ∞. Fixation of the cooperation allele becomes easier in this case as *π_A|a_* > 1/*N*_T_ translates to *D* > 2*C*. Conversely, fixation of the noncooperation allele also becomes easier when population structure is strong since the fixation condition becomes *D* < 2*C*. These conditions are summarized in Table 4.

**Table 4:**
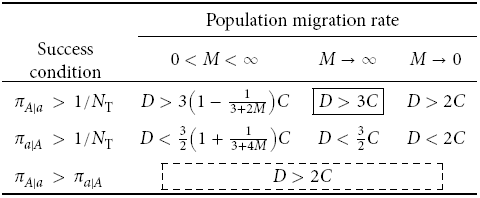
Evolutionary success conditions calculated in a infinite island model (*N* → ∞and *n*→ ∞) with hard selection. The solid boxed condition is the one-third law [116] and the dashed box condition is the risk dominance condition [34, 70, 78] that holds for all values of *M*.

| Success<br>condition | Population migration rate |  |  |
| --- | --- | --- | --- |
| | $0 < M < \infty$ | $M \rightarrow \infty$ | $M \rightarrow 0$ |
| $\pi_{A\vert a} > 1/N_T$ | $D > 3\left(1 - \frac{1}{3+2M}\right)C$ | $\boxed{D > 3C}$ | $D > 2C$ |
| $\pi_{a\vert A} > 1/N_T$ | $D < \frac{3}{2}\left(1 + \frac{1}{3+4M}\right)C$ | $D < \frac{3}{2}C$ | $D < 2C$ |
| $\pi_{A\vert a} > \pi_{a\vert A}$ | $\boxed{D > 2C}$ | | |

As discussed in section 2.4, each fixation condition alone (and, consequently, the one-third law, *π_A|a_* > 1/*N*_T_) is sufficient as a measure of evolutionary success only when genetic interactions are additive. When non-additive or synergistic interactions are included, the condition *π_A|a_* > *π_a|A_* (eq. 9) should be used. This condition is

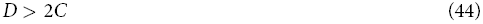

using the fixation probabilities in (41) and (42) (see Table 4). Interestingly, this implies that whether the cooperation allele *A* or the noncooperation allele *a* is more common at stationarity is independent of the strength of population structure, *M*, and depends only on the payoffs in the social game. This is a generalization of the Taylor cancellation result in the sense that the kind of simple population structure considered here (non-overlapping generations, homogenous island-type migration, hard selection) is not sufficient for the benefits of cooperation *B* to affect selection for cooperation. Rather, synergistic effects are important, but they must be significantly outweigh the costs in order for cooperation to be more prevalent than noncooperation.

In fact, once synergistic effects outweigh the costs at all, *D* > *C*, they change the structure of the social game from a Prisoner’s Dilemma where noncooperation is the strictly dominant strategy to a Stag-Hunt or coordination game where both cooperation and noncooperation are Nash equilibria [12]. In coordination games in unstructured populations, resident populations of cooperators and noncooperators are both resistant to invasion by the complementary type when evolution is deterministic (i.e., there is no genetic drift). This implies that whether allele *A* or *a* becomes fixed in the population depends on the initial frequency of *A*. When the initial frequency is greater than the mixed strategy equilibrium *z*\* = *C*/*D*, selection leads to fixation of the cooperation allele *A*, and fixation of the noncooperation allele *a* occurs for initial frequencies less than *C*/*D*. In effect, if the phenotypic space is the probability of cooperation *z*, then the mixed strategy equilibrium is a fitness valley and pure cooperation and noncooperation are fitness peaks [165]. The basin of attraction for the cooperation peak in this case would be (*z*\*, 1), and for the noncooperation peak the basin would be (0, *z*\*). An intuitive condition for the cooperation peak to be more likely to evolve is that is has the larger basin of attraction under a model of simple deterministic evolution. This condition, which is called “risk dominance” in game theory [70, 78], is equivalent to 1 − *z*\* > *z*\* or

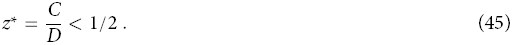

However, this is exactly the same condition we obtained from *π_A|a_* > *π_a|A_* in (44). The fact that we can derive the evolutionary success condition for cooperation in a structured population with selection, drift, and mutation from the risk dominance condition in a purely deterministic model with no population structure is another way of describing the generalized Taylor cancellation result introduced above.

Even though the equivalence between *π_A|a_* > *π_a|A_* and risk dominance is proven here for the infinite island model (*n* → ∞ and *N* → ∞), it approximately holds for the finite island model as well. In order to show this, we make two observations. First, examining the general condition for *π_A|a_* > *π_a|A_* in equation (24), we observe that so long as fitness depends on at most three way genetic interactions (*d* = 2), which is true for the fitness function in (39), all three-way coalescence times cancel out. Thus, we only need exact pairwise coalescence times and don’t need exact triplet coalescence times (we elaborate on further ramifications of this fact in the Discussion). Second, as suggested by Ladret and Lessard [86, p. 416], exact expected coalescence times can be calculated using a discrete-time Markov process that produces the same linear equations in Notohara [115, p. 66] that were used to generate the structured coalescent results in (40). These equations, given by (I) in Ladret and Lessard [86], produce the following exact expected coalescence times:

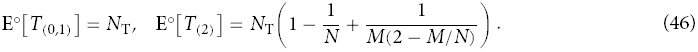

Combining the above expressions and the fitness function from (39) with the evolutionary success condition *π_A|a_* > *π_a|A_* in equation (24), we get

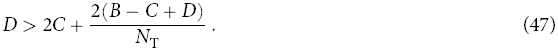

Condition (47) contains a single correction to the risk dominance condition, 2(*B* - *C* + *D*)/(*N*_T_), which is due to local competition in a finite population. Compared to the infinite island model, the cooperation allele *A* is only slightly less prevalent in this case. If the number of groups is infinite or if each group is infinitely large, condition (47) simplifies to risk dominance.

### 3.2. Demography and the scale of competition

Under the baseline demography of hard selection, we obtain the general Taylor cancellation result that removes the additive benefits of cooperation. This cancellation is a function of how demography shapes both the competitive environment and genetic identity within and between groups. Thus, demographies that create a different competitive environment may be more or less conducive towards the evolutionary success of the cooperation allele.

One of the most common alternative demographies to hard selection is soft selection [23] where individuals compete for resources or breeding spots within their group before the migration stage. Thus, each group contributes the same number of individuals to the next generation. Intuitively, this should make it more difficult for the cooperation allele *A* to succeed since groups with a higher frequency of *A* will not be more productive than groups with a lower frequency. In this case, the fitness function is

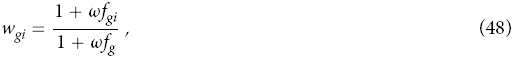

[132][p. 125] when the number of groups is large (*n* → ∞). For additive genetic interactions (*D* = 0) and using the exact pairwise expected coalescence times in (46), the evolutionary success condition *π_A|a_* > 1/*N*_T_ is

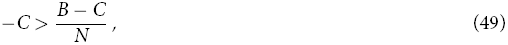

which agrees with previous analyses [96, 132]. In contrast to the case of hard selection where the evolutionary success condition is −*C* > 0, soft selection increases the strength of local competition so that cooperation is actually selected against in finite groups with a strength proportional to the net benefits, *B* - *C*. Allowing for non-additive interactions (*D* ≠ 0), we calculate the evolutionary success condition *π_A|a_* > *π_a|A_* to be

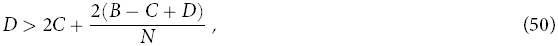

which again has a stronger local competition correction to the risk dominance condition than the hard selection case (eq. 47). In fact, since it doesn’t depend on the number of groups *n*, the soft selection correction is not simply a finite population size (*N*_T_ < ∞) effect and is due to competition occurring exclusively within groups.

A strong contrast to the soft selection demography is where the social interaction still occurs within the group but the scale of competition is the whole population. This might occur when groups directly compete for resources and successful groups produce propagules whose genotype frequency is proportional to their frequency within the group [e.g., 51, 94, 96]. If migration occurs at the adult stage, then the fitness function for this case is given by

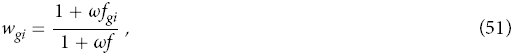

and the neutral demography is still represented by an island model whose expected coalescence times are given in equations (40). The evolutionary success condition *π_A|a_* > *π_a|A_* for this case is

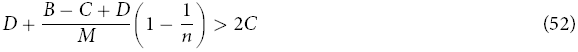

for large groups (*N* → ∞). When migration is high and the population is unstructured (*M* → ∞), we recover the risk dominance condition from (52). Strong population structure from weak migration however very strongly selects for cooperation. In fact, for any level of cost, there is a low enough population migration rate *M* that results in more cooperation alleles *A* at stationarity. This strong selection for cooperation is a direct result of the lack of local competition, which allows the benefits of cooperation to accrue to individuals who cooperate with other individuals in their group who tend to share their genes IBD.

## 4. THEORY: LONG-TERM EVOLUTION

We showed above how the forces of selection, mutation, and genetic drift generate the stationary distribution between two alleles when mutation is weak and evolutionary change follows the TSS. This stationary distribution gave us a measure of evolutionary success of one allele relative to another, which was simply *π_A|a_* > *π_a|A_*. Under the additional assumption of weak selection, we showed how to calculate this condition (eq. 24) for arbitrary non-additive genetic interactions. From this condition, we obtained Hamilton’s rule, the one-third law, risk dominance, and generalizations of these conditions. However, this condition only gives the stationary distribution among a fixed set of alleles and thus does not explicitly make predictions for longer timescales when the evolutionary process can sample a continuum of alleles.

Studying long-term evolution among a continuum of possible alleles requires specifying how those alleles are generated by mutation and how mutations are fixed or lost over the long-term due to selection and drift. Just as with the short-term model among a finite set of alleles, we will assume weak mutation so that the population can be described by the TSS and we only need to track the evolution of a population from one fixed, or monomorphic, state to another. Our approach to modeling the long-term process uses the “substitution rate” approach of Lehmann [88, 165], which derives from population genetic approaches to adaptation [55, 56] and to kin selection in finite populations [132, 134, 154, 158] and from the adaptive dynamics approach [19, 33, 107].

### 4.1. Substitution rate approach and the TSS diffusion

The essence of this approach is that, over the long term, the evolutionary process at each point in time can be fully characterized by a substitution or transition rate that measures how likely the population is to move from one monomorphic state to another. We assume that the substitution rate is an instantaneous measure of change, which is justified since organismal life cycles and generation times become very short on the scale of long-term evolution. If *ρ*(*z*_1_, *t*_1_|*z*_0_, *t*_0_) is the probability density that a population is monomorphic for trait *z*_1_ at time *t*_1_ given it was monomorphic for *z*_*o*_ at time *t*_0_, then we define the substitution rate as

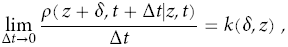

which is intuitively the rate at which mutations of type *z* + *δ* are produced and fixed in the population of type *z*. The substitution rate is a function of both the mutation rate, *μ*, and the distribution of mutational effects, *u*(*δ*, *z*), which represents the probability density that a mutant offspring is of type *z* + *δ* given that its parent is of type *z*. For simplicity, we assume that *μ* does not depend on the resident trait in the population, *z*. Using weak mutation and the TSS condition in (4), Champagnat [18, 21] showed that the long-term TSS can be characterized as a Markov jump process (continuous in time) with instantaneous jump rate

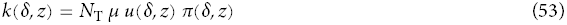

(see also Theorem 1.1 in [21] and eq. 2 in [88]). This rate is conceptually analogous to the classic long-term neutral substitution rate *k* = *N*_T_*μ*(1/*N*_T_) = *μ* from molecular evolution [83].

Following standard methods in stochastic processes, the Markov jump process representing the TSS with jump rate *k*(*δ*, *z*) can be represented with the following (forward) master equation [33, 49, 88]

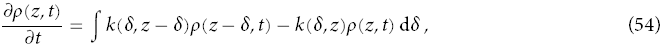

where we write *ρ*(*z*, *t*) = *ρ*(*z*, *t|z*_0_, *t*_0_) for simplicity. The master equation captures the intuition that the change in probability density for trait *z* at time *t* is equal to the sum of the jumps towards that trait minus the sum of the jumps away from that trait. Without further assumptions about the size of jumps, the master equation is difficult to analyze. A common way to approximate Markov jump processes is to assume that the jumps are small, which turns the discontinuous jump process into a continuous process and turns the master equation into a diffusion equation. Specifically, standard methods (e.g., a Kramers-Moyal expansion [49]) generate a diffusion equation by ignoring third-order and higher moments of the jump process. Biologically, we can justify this approximation by assuming that the mutational effects *δ* cluster tightly enough around the mean trait value *z* so that the evolutionary dynamic is affected only by the variance of the distribution of mutational effects and higher order moments can be neglected. The (forward) diffusion equation obtained with these methods for the jump process in (54) is

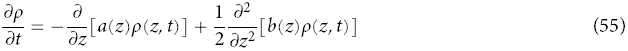

where

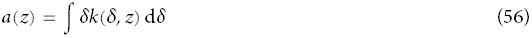

is the “drift” term and measures the mean jump away from the trait *z* and

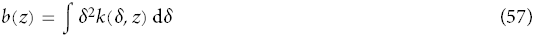

is the “diffusion” term and measures the variance of the jumps away from *z*. The drift and diffusion terms can be simplified by assuming that the fixation probability is differentiable (which is true if fitness is differentiable; see section 2.5) and approximating the substitution or jump rate by the first-order Taylor series

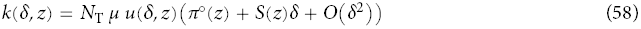

where *π*°(*z*) = *π*(0, *z*) and 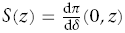 is the selection gradient. Using the expansion in (58) and assuming that the mutational distribution is symmetric in *Δ*, the drift and diffusion terms become, respectively,

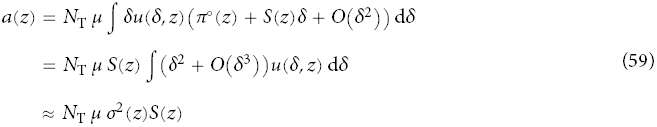

and

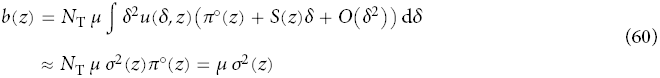

where *π*°(*z*) = 1/*N*_T_, *σ*^2^(*z*) = ∫ *δ*^2^*u*(*δ*, *z*) d*δ* is the second moment (and variance) of the mutational effects distribution and third-order and higher moments are neglected. Substituting these expressions for *a*(*z*) and *b*(*z*) into the diffusion equation (57) yields

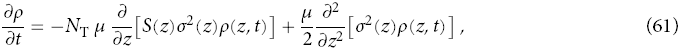

which was derived by Lehmann [88, eq. 4] and is analogous to the stochastic differential equation derived by Champagnat and Lambert [21, eq. 3]. The diffusion equation in (61) is our main mathematical description of how the TSS evolves over the long-term. In a sense, this diffusion equation is a stochastic version of the deterministic canonical equation of adaptive dynamics [33]. The first term in (61) measures the deterministic effect of selection on the trait and is the counterpart of the canonical equation of adaptive dynamics [21]. The second term measures the stochastic effect of genetic drift through the neutral fixation rate (eq. 60).

### 4.2. Evolutionary success in the long-term TSS

Just as in the short-term TSS analysis, we will define the evolutionary success for a trait *z* in the long-term TSS as its stationary probability or *ρ*(*z*) = lim_*t→∞*_ *ρ*(*z*, *t*). Using the long-term TSS diffusion in (61) and standard methods [41, 49, 81], we find that

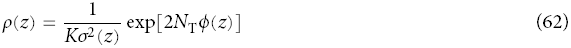

where

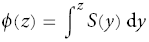

is the “potential” function and *K* is a normalizing constant that ensures *ρ*(*z*) integrates to one over its support. The most successful traits will be those that reside at peaks of the stationary distribution, and the least successful will reside at troughs. Obtaining the peaks and troughs, when they do not reside at the boundaries of trait space, requires calculating the extrema of *ρ*(*z*), which must satisfy

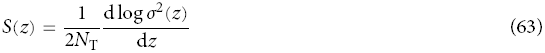

evaluated at a candidate extremum *z* = *z*\*. So long as either population size *N*_T_ is very large or the mutational variance *σ*^2^(*z*) does not depend on the resident trait *z*, equation (63) becomes

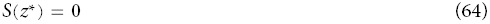

Consequently, in these two cases, the extrema of the stationary distribution are given the zeros of the selection gradient *S*(*z*), which are also the extrema of the fixation probability *π*(*Δ*, *z*) with respect to *Δ*. For the remainder of this analysis, we will assume that d*σ*^2^(*z*)/ d*z* = 0 and the extrema of the stationary density are zeros of the selection gradient.

The zeros of the selection gradient are the candidate evolutionary equilibria obtained using evolutionary game theory, adaptive dynamics, and inclusive fitness theory. A candidate evolutionary equilibrium *z*\* is called “convergence stable” when *S*′(*z*\*) < 0 [35, 37, 150], which means that for resident trait values close to *z*\*, mu-tants invade such resident populations only when those mutants are closer to *z* than the resident. Convergence stability is a natural way to characterize long-term evolutionary attractors (which may or may not be “branching points”; see refs [167] for the relevance of branching in the TSS). The condition for an extremum *z*\* of the stationary density *ρ*(*z*) to be a local maximum, d^2^*ρ*/ d*z*^2^ < 0, turns out to be precisely the convergence stability condition. Convergence stable traits may also reside on the boundaries of the trait space. In this case, the lower boundary is convergence stable when *S*(*z*) < 0 and the upper boundary is convergence stable when *S*(*z*) > 0. These two conditions correspond, respectively, to boundary maxima for the stationary density, d*ρ*/ d*z* < 0 for the lower boundary and d*ρ*/ d*z* > 0 for the upper boundary. Thus, long-term evolutionary attractors given by convergence-stable equilibria obtained from the selection gradient are generally local maxima of the stationary density of the long-term TSS diffusion [88].

In the simple TSS model above, the selection gradient also determines more generally which phenotypes are more probable than others at stationarity. This is a consequence of our assumption that both total population size *N*_T_ and mutational variance *σ*^2^ are not functions of the resident trait *z*. To see this, consider the relative likelihood of two traits, *z*_*A*_ and *z*_*a*_, at the stationary distribution: trait *z*_*A*_ is more likely that trait *z*_*a*_ when

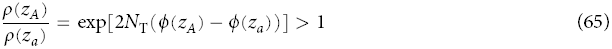

or

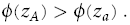

Thus, though total population size and mutational variance may affect how divergent the likelihoods of two traits are at stationarity, the selection gradient alone (through its integral) completely determines which trait is more likely than the other.

### 4.3. Additive genetic interactions and weak versus strong payoffs in the long-term TSS

Since the diffusion approach to the long-term TSS uses *δ*-weak selection (see equation 58), we know from section 2.5 that this approach can only capture additive genetic interactions with respect to fitness. This could suggest that the diffusion equation for the long-term TSS only represents evolution under additive genetic interactions. Intriguingly, this is not true as we will show in section 5 where we apply the long-term TSS diffusion to the same social game analyzed using the short-term process in section 3. Assuming that the payoffs to the social game are small (“weak payoffs”) and taking a large total population size limit (*N*_T_ → ∞), we find that the diffusion model of the long-term TSS can reproduce all the results of the short-term TSS with three-way genetic interactions in a group structured population. This is likely due to the cancelation of three-way coalescence times in short-term model.

Moreover, we find for”strong payoffs” that the diffusion approach appears to reproduce some results generated from another evolutionary game theory analysis that does not assume weak selection. Specifically, Fudenberg et al. [48] show that the condition for evolutionary success in an unstructured population is less favorable to cooperation than simple risk dominance would suggest, and the long-term TSS diffusion reproduces this effect.

## 5. APPLICATION: SOCIAL GAMES IN STRUCTURED POPULATIONS

In this section, we apply the long-term TSS diffusion to the same social game as in section 3 with a similar group-structured population. Our goal is to track the evolution of the continuous trait *z* that measures the fraction of time that an individual cooperates with social partners living in its own group; the complementary fraction 1 − *z* is the fraction of time the individual does not cooperate. Using the payoffs from Table 3, the fertility of individual *i* in group *g* is 1 + *f*_*gi*_ where

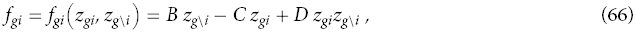

which is analogous to the fertility function in equation (37). Following the weak effect mutation model for the short-term TSS, we assume that the phenotype of individual *i* in group *g* is *z*_*gi*_ = *z* + *p*_*gi*_*δ* where *p*_*gi*_ is the frequency of the mutant allele in that individual.

### 5.1. Selection gradient

Without explicitly specifying how fertility and survival translate into fitness, we can calculate the selection gradient by assuming a homogeneous group-structured population where dispersal is potentially local so that genetic identity or relatedness can build up. This is the same kind of demography we used in section 2.7 where we reproduced Hamilton’s rule from the selection gradient in equation (34). Briefly, this demography ensures that the derivatives of fitness with respect to the trait values of different individuals at neutrality take only three possible values: 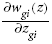 for the effect of the focal individual’s trait on its own fitness, 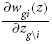for the effect of the average group trait (excluding the focal) on the focal’s fitness, and 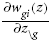 for the effect of the average trait in other groups (i.e., all groups except *g*) on the focal’s fitness. The latter derivative, 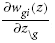, can be written in terms of the former two since the selection coefficients must sum to zero. The IBD probabilities also collapse to three categories: identity with self, which is one, identity with another individual in the group or *Q*_0_, and identity with an individual in another group or *Q*_1_. These facts together allow us to rewrite the selection gradient in equation (34) as

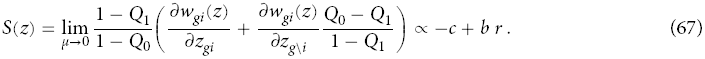

where we have used the fact that 1 − *Q*_0_ = 2*N*_T_*μ* + *O*(*μ*2) (eqs. 26 and 46 in [111] and eq. 3.68 in [132]). The selection gradient in (67) is the same as that derived by Rousset [132, 134] for group structured populations. Using the definitions for terms in Hamilton’s rule in section 2.7 where 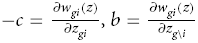 and *r* = *F*_ST_ = 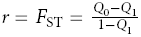, it is clear that selection gradient *S*(*z*) is proportional to the inclusive fitness effect, which is a function of the phenotypic trait *z* and all of the demographic effects that shape *b*, *c*, and *r*.

Since fitness is a function of fertility, we can write the selection gradient *S*(*z*) in equation (67) in terms of derivatives of fertility instead of fitness. This will allows us to express *S*(*z*) in terms of the payoffs *B*, *C*, and *D* from the social interaction. First, we assume a group-structured demography where fitness is determined by competition with groups for the *N* available spots or patches in the next generation. Let *α* be the probability than an offspring will compete for a patch in its natal deme and 1 − *α* be the probability that it competes in some other deme. The fitness of a focal individual *i* in group *g* is then

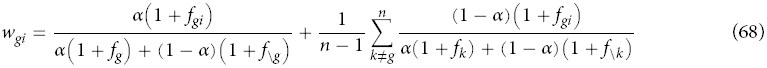

where 1 + *f*_g\*i*_ is the average fertility in group *g* excluding individual *i* and *f*_\*g*_ is the average fertility among all groups excluding *g*. Ignoring terms *O*(*δ*^2^) due to the assumption of *δ*-weak selection allows the average fertilities in the denominator of (68) to be rewritten as the fertility of an individual with the average phenotype:

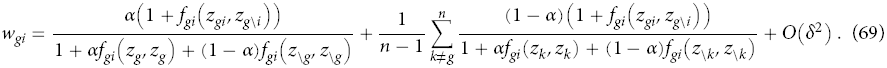

Using equation (69), we can write the fitness derivatives in the selection gradient in equation (67) as

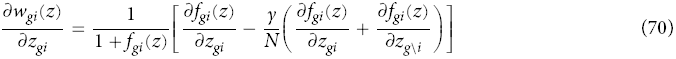

and

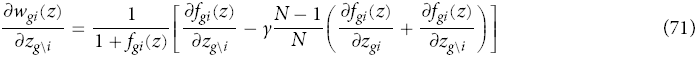

where

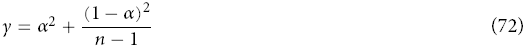

is the probability that two offspring will compete in the same group and is a measure of the strength of local competition (also called the “scale of competition” [44, p. 114]). Using the fitness derivatives in (70) and (71) in equation (67) allows us to rewrite the selection gradient as

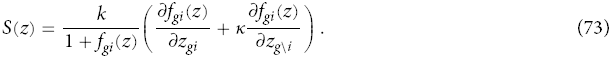

where

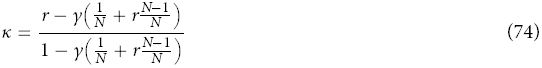

is the “scaled relatedness coefficient” and

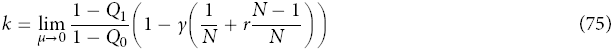

is a positive coefficient that scales the magnitude of the selection gradient and depends on demographic and competitive effects. We assume that both *κ* and *k* do not depend on the phenotype *z*, which is true if the demographic variables, such survival, migration, population size, etc, do not depend on *z*. Additionally, fertility must also be generally large or Poisson distributed in order to neglect the effect of the trait on demographic stochasticity [89].

The first term in the parentheses of (73) is the effect of the trait in the focal individual on its own fertility. The second term is the effect of the trait in the group (excluding the focal) on the fertility of the focal weighted by the scaled relatedness coefficient *κ*. The scaled relatedness in equation (74) is classic relatedness (*r*) reduced by the effect of local competition weighted by the average relatedness of the individuals affected by the local competition (*?*(1 + *r*(*N* - 1))/*N*) and normalized. Thus, scaled relatedness *κ* accounts for both relative genetic identity due to genetic relatedness and competitive effects due to demography and finite population size [96, 164, 165]. In general, the scaled relatedness *κ* can take a value between -1 and 1 depending on the demography [96]. For example, the hard selection demography in equation (38) has *α* = 1-*m* and from equation (74) yields *κ* = -1/(*N*_T_ - 1) [165, eq. B.1], soft selection in equation (48) has *α* = 1 and yields *κ* = -1/(*N* - 1) [96, eq. A-8], and group competition from equation (51) has *α* = 1/*n* and yields *κ* = *r* when *n* → ∞.

Compared to the fitness effects and genetic identity formulation of the selection gradient in (67), the selection gradient in (73) partitions terms into fertility effects and scaled relatedness. The former partition is that used to define *b*, *c*, and *r* in Hamilton’s rule, which leads to the definitions for different social behaviors based on fitness effects in Table 2. In the latter partition, all the effects of demography and local competition are encompassed by scaled relatedness *κ* and the coefficient *k* and the effects of the phenotypic trait on the social interaction are isolated in the fertility effects. In so far as we are interested in understanding how the immediate payoffs from social interactions and demography *independently* contribute to selection on social behavior, the latter partition is advantageous. Previous models have used this partition where *κ* is also called scaled relatedness [2, 96, 164] or “compensated relatedness” [61],”potential for altruism” [50], “potential for helping” [130], “index of assortativity *σ*_0_” [4], or the “structure coefficient *σ* “ [5, 146, 148]. Of particular relevance here, the structure coefficient *σ* introduced by Tarnita and collaborators [146, 148] has been used to analyze the evolution of discrete strategies in finite and structured populations where non-additive genetic interactions are possible due non-additive payoffs (i.e., *D* ≠ 0). As we will see below, our long-term TSS diffusion can reproduce a central result from the work using *σ* even though the selection gradient *S*(*z*) only captures additive genetic interactions at any one point in time.

Applying the fertility function in (66) to the selection gradient in (73) yields

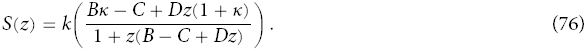

Recalling that *S*(*z*\*) = 0 define candidate evolutionary equilibria, we find one internal equilibrium at

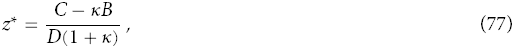

which is viable mixed strategy only when *D*(1 + *κ*) > *C* - *κB* > 0. When there is no effect of population structure and *κ* = 0, this simplifies to *D* > *C*, which implies that the social game is one of coordination where both the boundary equilibria *z* = 0 and *z* = 1 are convergence stable (i.e., *S*(0) < 0 and *S*(1) > 0) and the internal equilibrium *z*\* is unstable (*S*′(*z*\*) > 0). If *D* < *C*, then the game is Prisoner’s Dilemma and only *z* = 0 is stable. When population structure is at its strongest, *κ* = 1, *B* > *C* ensures that the internal equilibrium does not exist (not a valid mixed strategy) and only *z* = 1 is stable, which implies the social interaction is a mutualism game. Thus, non-additive payoffs, or synergy, can generate the possibility of cooperation by changing the game from a Prisoner’s Dilemma to a coordination game [2]. Population structure can also generate cooperation, but in a stronger way since when cooperation is stable, it is the only equilibrium.

### 5.2. Stationary distribution

Substituting the selection gradient in (76) into the stationary distribution in (62) yields

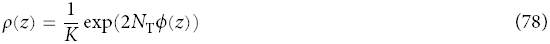

where

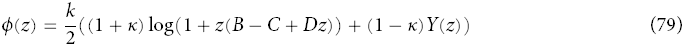

is the “potential function” [49],

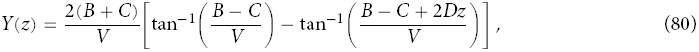

and 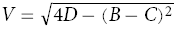. In the left panel of Figure 1, we plot the stationary density *ρ*(*z*) in (79) for *N*_T_ = 20, *B* = 1, *C* = 0.5, and *κ* = 0. As expected for *D* = 0 when the game is a Prisoner’s Dilemma, we see a peak in the density at *z* = 0 and the mean value of *z*, E[*z*], is close to *z* = 0. When *D* > *C* = 0.5, the social interaction becomes a coordination game and both full noncooperation, *z* = 0, and full cooperation, *z* = 1, become convergence stable. The full cooperation peak is initially very small, and the population spends most of its time with a trait value close to *z* = 0 until *D* increases to at least 2*C* = 1. Recall from condition (45) that *D* > 2*C* is the risk dominance condition that ensures the basin of attraction of full cooperation is larger than that of full noncooperation. The mean value of the trait E[*z*] crosses *z* = 1/2 at *D* ≈ 1.15 > 2*C*, which implies that cooperation under the long-term TSS model is more difficult to obtain than the risk dominance condition suggests. However, risk dominance appears to be the correct condition when the payoffs are much weaker. This is shown in the right panel of Figure 1 where *B* = 1 × 10^−2^, and *C* = 0.5 × 10^−2^. The lower values of the payoffs induce much weaker selection, which leads to a mean trait value much closer to 1/2 for all values of *D*, but the crossing point is almost exactly at *D* = 2*C* as risk dominance predicts. In fact, we will show that the risk dominance condition can be recovered from the stationary density *ρ*(*z*) as part of a more general condition for evolutionary success under weak payoffs.

**Figure 1:**
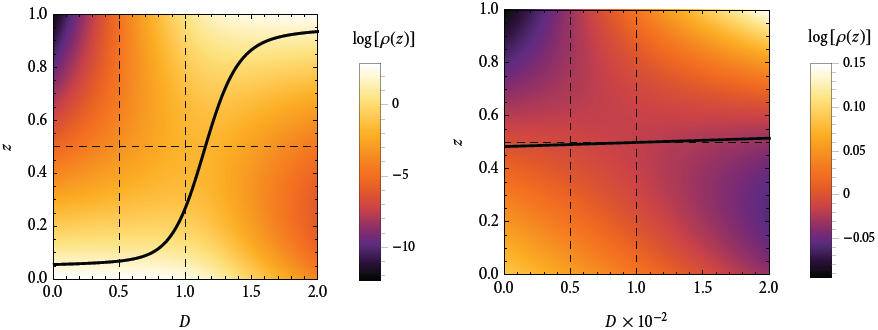
Log stationary density, log[*ρ*(*z*)], for *N*_T_ = 20 and *κ* = 0 plotted as a function of the synergy *D*×10 -2 parameter *D*. The solid black line represents the mean trait value, E[z]. The first vertical dashed line represents the boundary between a Prisoner’s Dilemma for *D* < *C* and a coordination game *D* > *C*. For *D* < 2*C*, full noncooperation (*z* = 0) is risk dominant (see condition -45and full cooperation is risk dominant for D > 2C.Left panel: B = 1, and C = 0.5. Right panel: B = 1 × 10^−2^, and C = 0.5 × 10^-2^.

The scaled relatedness *κ* has a strong effect on the stationary distribution of trait values [165], which we show in Figure 2 by plotting the mean trait value E[*z*]. In the upper left panel, the parameters are the same as Figure 1 except we vary *κ* from -0.5 to 0.5. When population structure increases genetic relatedness and does not induce much local competition, *κ* takes positive values. The plot shows that positive values of *κ* produce a shift in the E[*z*] curve so that lower values of the synergistic payoff *D* are required to obtain a high level of cooperation in the population compared to *κ* = 0 when there is no effect of population structure. The converse is true for negative values of *κ*, which require higher levels of *D* for any significant amount of cooperation. In these cases, IBD within groups is low and local competition between relatives is strong. These results are directly analogous to those obtained from the short-term TSS model in section 3. Instead of varying population structure continuously in the short-term TSS model, we analyzed different demographic scenarios and found that strong local competition led to a more stringent condition for the cooperation allele to be evolutionarily successful (eq. 50) and no local competition lead to a more relaxed condition (eq. 52).

**Figure 2:**
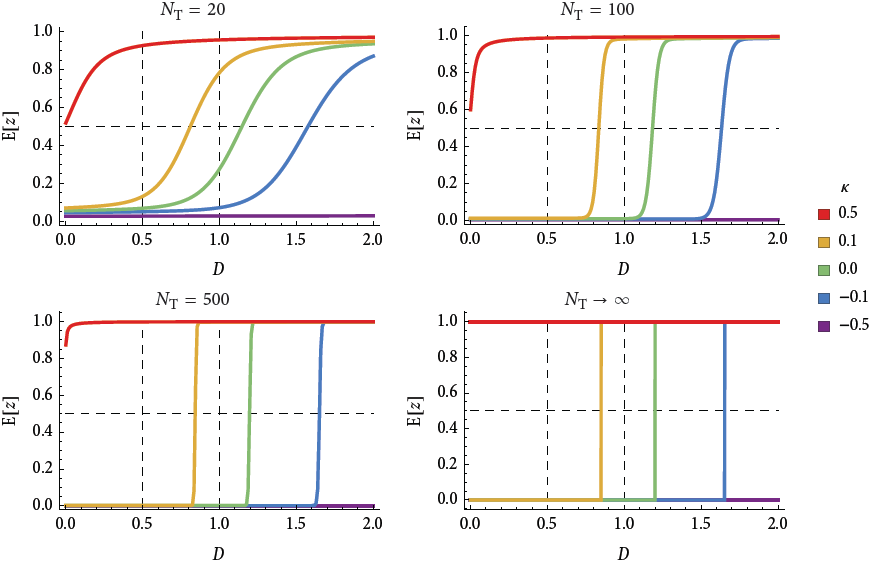
Expected trait value E *z* at stationarity under the long-term TSS diffusion (eq. 78) as a function of the synergistic payoff *D*. Total population size is given above each plot and scaled relatedness coefficients are given by the line colors in the legend varying from *κ* 0 5 in purple to *κ* 0 5 in red. The remaining payoffs are set at *B* = 1 and *C* = 0.5, which implies the game is a Prisoner’s Dilemma for *D* < 0.5 and a coordination game for *D* > 0.5. For *D* < 1, the trait *z* = 0, full noncooperation, is risk dominant, and full cooperation or *z* = 1 is risk dominant for *D* > 1.

### 5.3. Stochastic stability

The remaining panels in Figure 2 demonstrated the effect of increasing total population size *N*_T_. As *N*_T_ grows, the E[*z*] curves approach step functions that appear in the *N*_T_ → ∞ panel; for values of *D* below a threshold, E[*z*] = 0 and the population is fully noncooperative, and above the threshold, E[*z*] = 1 and the population is fully cooperative. When the total population size is small, genetic drift is strong and the peaks in the stationary density are small. This implies significant probability density for trait values distant from the peaks (see the right panel in Figure 1 where drift is strong relative to selection). Using the metaphor of a fitness valley, the population frequently crosses the valleys between small peaks due to small *N*_T_. Increasing *N*_T_ reduces the effect of genetic drift and increases the height of the peaks in the stationary density, which means most of the probability density is located at the peaks. In this case, the population rarely crosses fitness valleys. In the limit as *N*_T_ → ∞, the peaks of the stationary density get infinitely tall and the population only visits the peaks. The amount of time the population spends in fitness valleys goes to zero but crossings still occur frequently “enough” so that the population can escape a lower peak and visit a higher peak. Thus, only a small set of peaks are visited by the population in the long-run. Such peaks are called “stochastically stable states” and were first studied in game theory in models of agents with simple learning rules [13, 43, 78, 118, 140]. A trait *z* is stochastically stable when

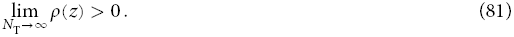

Van Cleve and Lehmann [165] prove that only the highest peak, as measured by the potential function *ϕ*(*z*) whose peaks are those of *ρ*(*z*), is stochastically stable. Since peaks in the stationary density correspond to convergence stable equilibria, only the convergence stable equilibrium associated with the highest peak is stochastically stable. In our current model of social interaction in a group structured population, the lower right panel of Figure 2 shows the stochastically stable value of *z* as a function of *D* and *κ*.

The advantage of identifying the stochastically stable state is that it is often unique as a function of the demography, *κ*, and the payoffs of the social game, *B*, *C*, and *D*. Using the stationary density in (78), we can show [165] that full cooperation (*z* = 1) is stochastically stable when

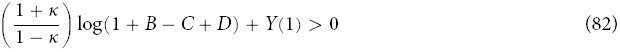

and full noncooperation (*z* = 0) is stochastically stable when the opposite condition holds. Condition (82) reveals immediately the positive effect that population structure has on the stochastic stability of cooperation since the left hand side is an increasing function of *κ*. Holding *κ* constant however, the relationship between condition (82) and the risk dominance condition is still difficult to discern. The results in Figure 1 suggest that stochastic stability in the long-term TSS diffusion and risk dominance may coincide when the payoffs are weak. Figure 3 shows more evidence for this by plotting E[*z*] for the same values of *N*_T_ and *κ* as Figure 2 except the payoffs are multiplied by 10^−2^. As population grows large, the *κ* = 0 curve crosses E[*z*] = 1/2 at exactly the risk dominance prediction of *D* = 2*C* = 1. In fact, we can show this analytically by assuming that *B*, *C*, and *D* are small and ignoring quadratic and higher terms in condition (82), which produces

**Figure 3:**
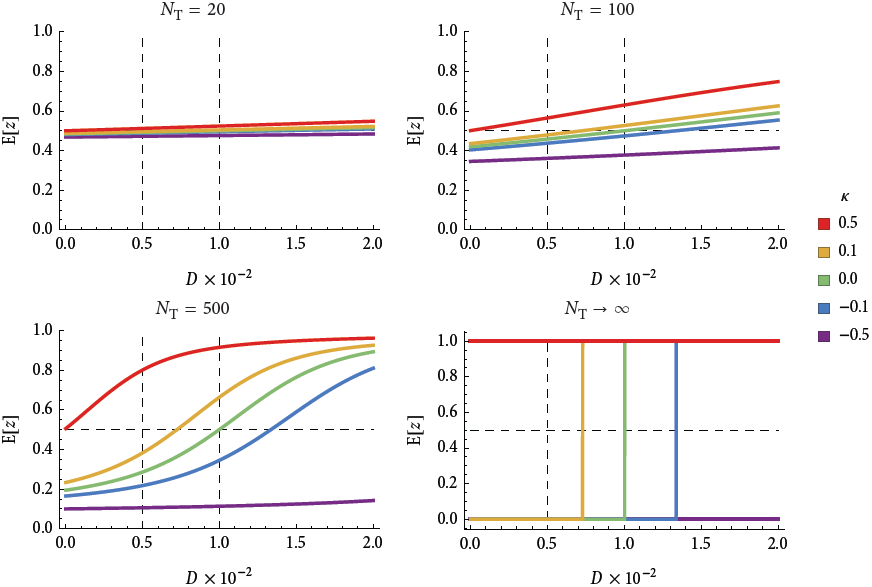
Expected trait value E[*z*] at stationarity under the long-term TSS diffusion as a function of *D* for weak payoffs where all payoffs are scaled by 10^−2^. Plots are otherwise identical to Figure 2.

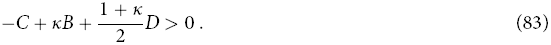

This is a “weak payoff “ stochastic stability condition. It is exactly the risk dominance condition when one uses the mixed strategy from equation (77) in the condition *z*\* < 1/2.

The weak payoff stochastic stability condition in (83) matches exactly with the results derived using the short-term TSS. For example, when *κ* = 0 and there is no effect of population structure, condition (83) simplifies to the risk dominance condition *D* > 2*C* derived in the short-term analysis by calculating *π_A|a_* > *π_a|A_* under hard selection. The scaled relatedness is *κ* = 0 for hard selection since local competition exactly cancels the benefits of cooperating with relatives [96, eq. 3.8]. In fact, we can rederive other results from the short-term TSS model by inserting appropriate values of *κ* into equation (83). For example, using *κ* = -1/(*N* - 1) for the soft selection demography in condition (83) reproduces condition (50). The scaled relatedness for the group competition demography is *κ* = *r* = *F*_ST_ under the infinite island model, which yields *κ* = 1/(1 + 2*M*) [96, limit as *Nm* → *M* and *N* → ∞ of eq. A-3], and this value of *κ* reproduces condition (52) when the number of groups is infinite. These results are summarized in Table 5. Additionally, condition (83) is the same as the one derived by Tarnita et al. [146, eq. 4] if one exchanges our scaled relatedness *κ* for their structure coefficient *σ* (specifically, *σ* = (1+ *κ*)/(1-*κ*)). It is notable that the condition of Tarnita et al. [146] is derived from a discrete strategy model under weak mutation and weak selection that is analogous to our short-term TSS whereas our long-term TSS diffusion result is obtained under *δ*-weak selection and weak payoffs.

**Table 5:**
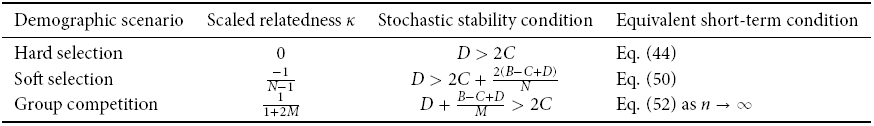
Long-term TSS stochastic stability condition with weak payoffs (equation 83) with different values of scaled relatedness *κ* for different demographic scenarios.

| Demographic scenario | Scaled relatedness $\kappa$ | Stochastic stability condition | Equivalent short-term condition |
| --- | --- | --- | --- |
| Hard selection | 0 | $D > 2C$ | Eq. (44) |
| Soft selection | $\frac{-1}{N-1}$ | $D > 2C + \frac{2(B-C+D)}{N}$ | Eq. (50) |
| Group competition | $\frac{1}{1+2M}$ | $D + \frac{B-C+D}{M} > 2C$ | Eq. (52) as $n \rightarrow \infty$ |

The ability of the weak payoff stochastic stability condition of the long-term TSS to reproduce the weak selection results for the short-term TSS suggests that strong payoffs might reproduce strong selection results from the short-term TSS. Our evidence for this conjecture is less conclusive since there is no known approximation of the evolutionary success condition *π_A|a_* > *π_a|A_* under strong selection and generic population structure. However, we can gain some intuition from a result from Fudenberg et al. [48] who analyze social interactions in a finite Moran model with no population structure. Assuming that *N*_T_ = *N* → ∞ and that selection strength is arbitrary (*ω* = 1 and arbitrary *B*, *C*, and *D*) in the short-term TSS, they find that *π_A|a_* > *π_a|A_* when

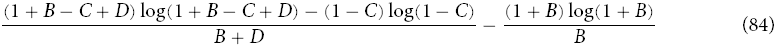

[48, Theorem 2 part b.3]. If we assume weak selection in condition (84) by replacing each parameter with its value times the selection strength *ω* and ignoring terms *O*(*ω*2), we recover the risk dominance condition in (45). For strong selection, condition (84) and the long-term TSS stochastic stability condition (83) are not equivalent but yield similar numerical results. Evidence of this match is in Figure 4, which plots the value of *D* at which the stochastic stability condition (83) is an equality as a function of scaled relatedness *κ*. The black curve is the strong payoff case (*ω* = 1) and the blue and red curves are weak payoffs (*ω* = 10^−2^ in blue and *ω* = 10-4 in red). The equivalent point from condition (84) where *κ* = 0 is plotted with the circled dots using equivalent selection strengths plotted in the same colors. It is evident from the figure that the short-term TSS results from Fudenberg et al. match very closely to the stochastic stability results from the long-term TSS diffusion. Both results show that increased selection strength requires a higher value of synergy *D* for full cooperation to be evolutionary successful than suggested by risk dominance. The correspondence between the short-term TSS model under weak selection and the long-term TSS diffusion under weak payoffs and their numerical match under strong selection and strong payoffs, respectively, suggests that the long-term process might capture the full potential of selection to shape social interactions under the TSS.

**Figure 4:**
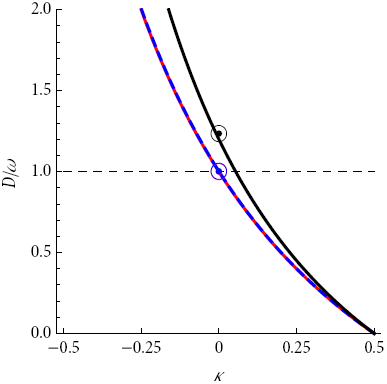
Threshold value of *D* according to condition (82) above which *z* = 1 and below which *z* = 0 are stochastically stable (*N*_T_ → ∞) as a function of scaled relatedness *κ*. Payoffs are scaled by selection intensity *ω* where the black curve has *ω* = 1, dashed blue *ω* = 10 ^–2^, and red *ω* = 10^−4^. The circled dots represent the analogous threshold calculated for a panmictic population using condition (84) from Fudenberg et al. [48]. Payoffs are set at *B*=1 and *C*=0 5.

## 6. DISCUSSION

Due in part to its simplicity, Hamilton’s rule has proved to be a remarkably useful tool for deriving insight into the effect of natural selection on the evolution of social behavior. By emphasizing the role of genetic correlations between individuals, Hamilton’s rule catalyzed interest in the effect of genetic population structure on all kinds of social behavior from parental care and cooperative breeding to cooperative hunting and colony defense. Hamilton’s rule also suggested a simple way to categorize social behaviors based on how they affect the fitness of focal actors and social partners (Table 2).

Early theoretical investigations of Hamilton’s rule and inclusive fitness quickly identified the source of this simplicity in two key assumptions of the rule, namely weak selection and additive genetic interactions with respect to fitness. The general utility and limitations of these assumptions for populations with arbitrary group or class structure became clearer in approaches that use an individually centered approach like the Price equation that describes evolutionary change via the statistical moments of the allele frequency distribution. Such approaches usually assume weak selection, which allows the calculation of higher-order moments via a quasi-equilibrium (QE) approximation. For the case of evolution at a single locus in a group-structured population, the QE approximation entails simply calculating *F*_ST_ or some other measure of population structure under neutral evolution. Assuming genetic additivity in addition to weak selection guarantees that Hamilton’s rule is independent of allele frequency [134], which means that it predicts both invasion and fixation of a mutant allele in a monomorphic population.

### 6.1. The trait substitution sequence and short and long-term evolution

Even with weak selection, polymorphisms are still possible since non-additive interaction can generate stabilizing selection and high mutation rates can maintain heterozygosity. If mutation rates are low enough relative to population size so that fixation or extinction occurs more quickly than a mutation, then only a single mutation will segregate in the population. This low mutation assumption, given in condition (4), generates the trait substitution sequence (TSS) whose short and long-term dynamics can be described in a remarkably complete fashion. Using weak selection and weak mutation, we showed above how to analyze evolutionary change under the TSS both in the short term when the set of possible alleles is finite (e.g. single nucleotide polymorphisms) and in the long term when a continuum of alleles is possible (e.g. morphological traits shaped by multiple cis-regulatory elements).

This difference between short versus long-term evolution with respect to the TSS is related to the broader conception of short and long-term evolution by Eshel [36, 37], Hammerstein [68, 69, “streetcar theory”], and others [173]. Growing out of an attempt to reconcile explicit population genetics approaches with phenotypic approaches from evolutionary game theory, Eshel characterizes short-term evolution as where, within a set of fixed genotypes, “natural selection will operate, in the short run, to change the genotype frequencies toward a new internally stable equilibrium” [37, p. 489]. Such an equilibrium can be monomorphic or polymorphic due to any process that could generate stabilizing selection such as heterosis, epistasis, or local adaptation in a structured population. Eshel defines long-term evolution as “characterized by the repeated introduction of new mutations into the population and in between periods of changes of genotype frequencies (say, short-term evolution) within the new simplex of genotypes” [37, p. 489]. Determining whether or not a new mutation will invade a generic internally-stable equilibrium is very difficult though extremely suggestive results were obtained by Eshel and Feldman [38], Liberman [101], and Hammerstein and Selten [69]; broadly, these authors found that new mutations cannot invade when the internally-stable equilibrium generates an ESS phenotype (under a linear population game) and that invading mutants shift a population towards an ESS if the population is already within a neighborhood of the ESS [38, 101]. This is a more general result than those obtained from our long-term TSS in that there are no restrictions on mutation rate or selection strength, which allows for complex polymorphisms. However, the assumptions of the TSS allows us to characterize both the short and long-term dynamics completely, which is not possible in the more general case of long-term evolution.

### 6.2. The short-term TSS

Beginning with the short term TSS, we described how a natural condition for evolutionary success, the expected frequency of the mutant allele being greater than the resident or E[*p*] > 1/2, is equivalent under weak mutation to the fixation probability of the mutant being larger than the resident or *π_A|a_* > *π_a|A_* [7, 47, 134]. Under weak selection, fixation probabilities can be written in terms of selection coefficients and expected coalescence times (eq. 22), which yields the general expression for *π_A|a_* > *π_a|A_* given in (24). This general condition, which is new to the literature, reveals how non-additive genetic interactions quickly increase the sensitivity of evolutionary success to pairwise, triplet, and higher-order coalescence times. When genetic interactions are additive, the condition *π_A|a_* > *π_a|A_* is equivalent to the simpler condition that the derivative of the fixation probability, or selection gradient, be positive: 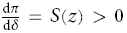. This latter condition readily reproduces Hamilton’s rule.

The appearance of expected coalescence times in the approximation of fixation probabilities [95, 100, 131] is useful both from conceptual and practical perspectives. Conceptually, expected coalescence times arise directly out of the fact that we calculate fixation probabilities after the generation of a new mutation in a monomorphic population and before another mutation enters the population [131]. Moving backwards in time, coalescence times express the effect of population structure and demography on the genealogy of the invading mutant. How the genealogy interacts with selection on the mutant depends on the fitness function; fitness functions linear in mutant allele frequency generate dependence on pairwise coalescence times, quadratic fitness functions generate coalescence times between three lineages, and so on. Practically, using expected coalescence times allow us to leverage the extensive results in coalescence theory [113, 170] that detail the effect of demography [e.g., 129] and population structure [e.g., 115, 168] on coalescence times.

Using Slatkin’s formula (eq. 33) for relating expected pairwise coalescence times to pairwise IBD probabilities [143], we showed how for additive genetic interactions the evolutionary success condition 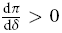 can be written in terms of IBD probabilities (eq. 34). This approximation in terms of IBD probabilities can also be obtained by calculating the evolutionary success condition

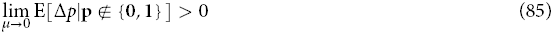

for additive genetic interactions [134]. Allen and Tarnita [7] showed that condition (85) is equivalent to *π_A|a_* > *π_a|A_*, which says that calculating condition (85) can yield the equivalent of condition (24) except with IBD probabilities instead of expected coalescence times. Our application of the Slatkin formula effectively allowed us to show this equivalency for additive genetic interactions; showing this equivalency for non-additive interactions requires relating coalescence times and IBD probabilities among an arbitrary number of lineages, which is a task for future work.

Using the success condition *π_A|a_* > *π_a|A_*, we reproduced and generalized several important previous results from group-structured (island-type) populations with the non-additive social interaction in Table 3. First, we reproduced the expressions for fixation probability *π_A|a_* given by Ladret and Lessard [86], which easily generate the one-third law of evolutionary game theory [116] by evaluating *π_A|a_* 1 *N*_T_ as population structure disappears or *M* → ∞. Applying the evolutionary success condition *π_A|a_ > π_a|A_* yielded the well-known risk dominance condition regardless of the level of population structure as measured by *M*. While risk dominance was initially proposed as a condition for determining which Nash equilibrium is optimal in two-player games in economics [70], it was subsequently shown to predict the stochastically stable strategy in population games where agents update their strategy by learning the best response [78] even when agents can only learn from local neighbors [14, 34]. Thus, these results confirm an important connection between the process of strategy selection in economic models and the process of natural selection with a certain type of population structure. Risk dominance also can be seen as a generalization of the classic cancellation result of Taylor [151, 152, 156] and others [128, 180] that says localized dispersal or population viscosity alone is not enough to create selection for cooperation or altruism. The cancellation results shows that the benefits of cooperation are exactly canceled by the costs of competing locally with kin when the population is homogeneously structured, density-dependent regulation occurs after dispersal (hard selection), and generations are non-overlapping. Our risk dominance result generalizes the cancellation result because it shows that the condition for cooperation to be evolutionarily successful is still independent of the amount of population structure even with non-additive or synergistic payoffs.

Moreover, our results go further because we showed how the risk dominance condition holds only in the limit of large total population size for the baseline demography of hard selection. When the total population size is small, the cancellation results no longer holds and local competition with kin degrades some of the benefit cooperators obtain when interacting with one another. More generally, the scale of competition can either increase or decrease the strength of local competition relative to relatedness, which makes selection for cooperation more or less stringent. Demographies with soft selection, where density-dependent regulation occurs before dispersal and competition occurs within the group, have greatly increased local competition that depends not on the total population size, but on the local group size *N*. In contrast, when the scale of competition is the total population and competition is effectively between groups, local competition is negligible and selection for cooperation depends primarily on the degree of genetic relatedness. If relatedness is strong enough (*M* is small), then the evolutionary success of the cooperation allele can be guaranteed. We can also relate these results back to the results in economics that suggest risk dominance is equivalent to stochastic stability in structured populations [14, 34, 78] (see also Sandholm [140] for a refinement of risk dominance). Our results suggest that models of learning and best-response dynamics in economics make strong implicit biological assumptions that eliminate the potential of population structure to affect strategic evolution; thus, these biological assumptions should be clearly specified and studied in economic models.

### 6.3. The long-term TSS

Moving to the long-term TSS and a continuum of possible traits, we applied a substitution rate approach that assumes the long-term process can be fully characterized by the rate a population jumps from one monomorphic state to another. This substitution or jump rate to a particular mutant trait *z*+ *δ* from a resident trait *z* is the number of mutants of type *z* + *δ* times the probability these mutations fix. The substation rate in the long-term TSS is analogous to the neural substitution rate of molecular evolution [83] except that the substitution rate is an explicit function of trait value, population size, and other parameters. By assuming that mutational effects are tightly clustered around the resident value, the substitution rates leads directly to a diffusion process [88] that characterizes trait change under the long-term TSS (eq. 61), which is a stochastic analogue of the canonical diffusion of adaptive dynamics [21].

From the long-term TSS diffusion, we obtained the stationary long-term trait density *ρ*(*z*) that can be used to asses evolutionary success. Specifically, local peaks in the stationary density correspond to classic evolutionary stable states [106] in linear or discrete games. In continuous trait games, these peaks correspond to convergence stable states [24, 35], which are attracting evolutionarily stable states where the selection gradient crosses zero from above [88, 165]. Being located at a peak in the stationary density is then a natural criterion for evolutionary success. Moreover, as the total population size goes to infinity and genetic drift becomes extremely weak compared to selection, only the highest peaks retain any positive probability at stationarity; all other peaks, even if locally convergence stable, are visited by the population with zero probability. The traits at these highest peaks are called stochastically stable. Since stochastically stable states are often unique, they represent one of the simplest possible predictions for trait evolution in the long term (a kind of “phenotypic gambit” [59]).

Using the stochastic stability criterion (eq. 81), we showed that the long-term TSS diffusion can a reproduce a range of results from the short-term TSS model presented here as well as results from other work using TSS-type assumptions. To show this, we first transformed the selection gradient from a Hamilton’s rule form with fitness effects and genetic relatedness to a form with fertility effects and “scaled relatedness” or *κ*. The scaled relatedness combines the effects of demography on both genetic identity and local competition whereas the fertility effects are strictly functions of the payoffs from the social interaction. Assuming “weak payoffs”, the condition for stochastic stability using this expression for the selection gradient matches exactly with the previous condition from Tarnita et al. [146] for the cooperation allele to be evolutionarily successful (*π_A|a_* > *π_a|A_*) where scaled relatedness *κ* corresponds to their structure coefficient *σ* = (1 + *κ*)/(1 − *κ*). Moreover, by using the values of scaled relatedness *κ* that correspond to the soft selection and group competition demographies, we reproduced the evolutionary success conditions derived from the short-term TSS. This is despite the fact that the selection gradient used in the long-term TSS depends only on additive genetic interactions, whereas the results from the short-term TSS and from Tarnita et al. [146] include non-additive interactions. We suggest below why the additive selection gradient can recover the results of the non-additive analyses. Finally, it is important to note that assuming weak payoffs does not imply that genetic drift dominate selection; since we assume that total population size *N*_T_ grows arbitrarily large, even small values of *B*, *C*, and *D* will eventually become important. These different notions of “weak” selection are described in section 6.5.

An intriguing aspect of the long-term TSS diffusion is that it only reproduced the short-term TSS results after assuming weak payoffs in the selection gradient. This suggested that strong payoffs in the selection gradient might produce results analogous to strong selection in the short-term TSS. In fact, we found some numerical evidence for this by comparing the results of the long-term TSS diffusion with the strong selection results of Fudenberg et al. [48] (see Figure 4) who derived a condition for stochastic stability in a panmictic population. Both results showed the same pattern, namely that the cooperative strategy was less likely under strong selection or strong payoffs than would be predicted by risk dominance, which is obtained with weak selection and payoffs. In the long-term TSS diffusion this effect can be explained by recalling the selection gradient *S*(*z*) from equation (76). In that expression, we can see that the fully cooperative strategy *z* = 1 in effect suffers a competitive cost relative to the noncooperative strategy *z* = 0 since the benefits of the cooperative strategy must be normalized by the mean benefit in the group. In contrast, assuming weak payoffs linearizes *S*(*z*) in terms of the payoffs *B*, *C*, and *D* and eliminates this cost.

A more general comparison between the long-term TSS diffusion and strong selection in structured populations remain to be done due to the lack of analytical results in that area (however see [110]) and the difficulty in defining a scaled relatedness *κ* for strong selection. However, even this preliminary evidence suggests that the long-term TSS may be a powerful tool for the analysis of social evolution even under some cases of strong selection.

### 6.4. Non-additivity and the power of the additive long-term approach

One of the crucial points of this article is that our derivation of Hamilton’s rule in the context of a dynamical model with selection, mutation, and genetic drift relies on three assumptions: (i) weak selection, (ii) weak mutation to ensure the TSS, and (iii) additive genetic interactions. The impact of breaking the last assumption and allowing non-additivity or synergy has been the subject of much interest [e.g., 52, 71, 93, 119, 127, 149, 157], and our analysis of the short-term TSS reinforces the importance of non-additivity in shaping the evolution of social outcomes. Nevertheless, our analysis of the long-term TSS also shows that even additive interactions, which generate Hamilton’s rule and the selection gradient used in the long-term TSS, can still generate useful and even powerful results. This is in contrast to some suggestions that analyses based on inclusive fitness methods are severely limited due to their assumption of additivity [6, 117], which does not appear to hold in some empirical systems [25, 58, 103]. In order to better understand precisely the extent to which additivity is limiting, we need to clarify the different levels at which additivity can hold and understand why additivity of genetic interactions is more powerful than commonly assumed.

Throughout this article, we have used additivity to refer specifically to additive genetic interactions with respect to fitness; this is equivalent to additivity of fitness effects with respect to allele frequency (*d* = 1 in eq. 17). With weak selection and the short-term TSS, we showed that additive genetic interactions lead directly to Hamilton’s rule since the selection gradient becomes a sum of selection coefficients times pairwise expected coalescence times or IBD probabilities. Analyzing the social interaction in Table 3 where individuals carry either a cooperation (*A*) or noncooperation allele (*a*), we showed how assuming additivity of payoffs (i.e., the synergistic payoff *D* = 0) implies additive fitness effects. However, the converse is not true; additive fitness effects do not imply that payoffs in the social interaction are additive.

To see this, consider an arbitrary fitness function for individual *i*, *w*_*i*_(**z**), which is a function of the (continuous) trait values of all other individuals in the population. The fitness function depends on the traits values of other individuals in part because individual *i* obtains payoff from a social interaction with some set of social partners. This payoff can be highly nonlinear as a function of the trait values of the social partners, which results in nonlinearity of the fitness function. However, so long as the fitness function is differentiable with respect to the traits values, a weak selection expansion of the fixation probability with respect to the mutant deviation *Δ* can still be calculated, and the resulting *Δ*-weak expansion has additive fitness effects. Thus, assuming a differentiable fitness function and *Δ*-weak selection imply that the selection gradient *S*(*z*) predicts the direction of selection and that Hamilton’s rule holds, even when payoffs are non-additive.

A simple example of a nonlinear payoff function is equation (66), which comes from the social interaction in Table 3. We showed this payoff function leads to the selection gradient in equation (76). We can apply the short-term TSS *δ*-weak condition, *S*(*z*) > 0, to this selection gradient and determine for any probability *z* of choosing the cooperation strategy whether a mutant trait *z* + *δ* is more common under the stationary distribution, even for a synergistic payoff *D* ≠ 0. In general, the nonlinearities of the payoff function will have important effects which traits are evolutionarily successful under the short-term TSS or convergence stable under the long-term TSS; for example, synergistic effects as measured by mixed partial derivatives 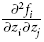 can have a powerful effect on the evolution of cooperative behavior [2, 3]. Such effects are important even in the absence of population structure [3] when such behavior would be a mutualism according to Hamilton’s rule (see Table 2). Cooperation is generated in these cases by reciprocity or behavioral responses [3], which can also enhance cooperation in structured populations where relatedness is important [2].

The weakness of the short-term TSS with respect to additive genetic interactions is that it cannot directly compare the relative evolutionary success of traits that differ by more than *δ*, such as full cooperation (*z* = 1) and full noncooperation (*z* = 0). Comparing the evolutionary success of full cooperation and no cooperation with the short-term TSS requires non-additive genetic interactions. In contrast, the long-term TSS measures the relative evolutionary success of full cooperation and noncooperation using the diffusion process to integrate population jumps over the whole trait space. Under weak payoffs for the long-term TSS, we showed that the diffusion approach produces the same results as the short-term TSS *even though the short-term TSS condition does not exclude non-additive genetic interactions and the long-term diffusion does*.

The likely explanation for this unexpected feature of the long-term TSS diffusion is that the social interaction we study (eqs. 37 and 66) has at most triplet genetic interactions and the triplet expected coalescence times cancel out of the short-term TSS condition (eq. 24); in other words, the short-term evolutionary success of cooperation versus noncooperation even with synergy (*D* ≠ 0) *depends only on pairwise measures of genetic identity*. This fact is easily missed when analyzing the conditions *π_A|a_* > 1/*N*_T_ and *π_a|A_* > 1/*N*_T_ independently, as is done in analyses based on the one-third law, since the triplet coalescence time terms cancel only in the full evolutionary success condition *π_A|a_* > *π_a|A_*. For more complex non-additive interactions, condition (24) reveals that higher-order expected coalescence times may not cancel, which suggests that the long-term TSS diffusion may have more difficulty approximating these interactions. The relationship between the long-term approach based on the selection gradient and arbitrary non-additive genetic interactions remains a topic ripe for further development. Nonetheless, the long-term TSS may be a powerful approach for modeling the evolution of social traits in structured populations under simple forms of non-additivity and potentially under strong selection.

### 6.5. The different “strengths” of selection in the TSS

In population genetics, “weak selection” typically means that selection is weaker than genetic drift in determining the fixation probability of an allele. The relationship between this notion of weak selection and the weak selection used in our study of both the short-term and long-term TSS deserves a bit of explantation. In both cases, the parameter controlling the strength of selection (*ω* for discrete phenotypes in the short term and *Δ* for continuous ones in the long term) is assumed to be small enough so that a separation of timescales occurs where IBD probabilities like *Q*_0_ and genetic associations like *F*_ST_ are well approximated by their values under neutrality. This means that the forces shaping the neutral values of these associations, mutation, migration, and genetic drift in the population models considered here, are strong relative to selection.

The TSS condition in (4) ensures that mutations are rare enough that they do not affect neutral genetic associations (i.e., we can calculate genetic associations using identity by descent instead of identity by state). Selection strength *δ* (for example) is weak relative to migration when *δ* ≪ *m* [137, 169]. Genetic drift in populations with restricted dispersal, such as those described by the island model, occurs within demes much more quickly than allele frequency change in the total population due to selection when deme size *N* and selection strength *Δ* are sufficiently small: *NΔ* ≪ 1 [22]. In this case, genetic associations like *F*_ST_ are well approximated by their neutral values. However, if the number of demes *n* is large enough, *nNδ* = *N*_T_*δ* > 1, which means that selection can still have a significant effect on the fixation probability compared to the neutral fixation probability of 1/*N*_T_. Thus, even though an assumption of *Δ*-weak selection (or *ω*-weak selection) allows genetic associations to be approximated by their neutral values, selection can still be strong enough to dominate the effect of genetic drift on the probability of fixation of a mutant allele in the total population so long as *N*_T_ is large enough.

In relation to the stationary distribution of the long-term TSS diffusion in equation (62), large *N*_T_ means that the relative likelihoods of different phenotypes grow divergent (see equation 65). In the limit as *N*_T_ → ∞, the stationary density becomes infinitely high at a single phenotype (or a small number of phenotypes), which is the stochastically stable state (see section 5.3). This is a kind of long-term version of strong selection where the stationary density is characterized entirely by the dynamics of selection through the selection gradient *S*(*z*). In contrast, when *N*_T_ is small, the stationary density approaches a uniform distribution, which is the outcome of the symmetric mutation distribution fixing mutations by genetic drift.

### 6.6. Limits of the TSS

While both the short and long-term approaches to the TSS have been powerful tools in understanding evolutionary forces on social behaviors, both approaches have their limits as does the TSS more generally. The long-term approach is specifically limited by its assumption of weak effect mutations, which follows from the *Δ*-weak selection assumption and implies genetic interactions must be additive. Though we show how this not a limitation for three-way interactions due to their cancelation in the short-term model, it is a limitation for higher-order interactions that are generic in *n*-player games. These interactions generate complex equilibrium structures [10, 57, 109, 123] and make *n*-player games particularly difficult to analyze in structured populations. Even though a *Δ*-weak approximation can be applied in such cases [124, 165], strong effect mutations (i.e., discrete phenotypes) are likely to have complex dynamics for *n*-player games that cannot be represented without an explicit consideration of higher-order genetic interactions.

Assumptions of both the short and long-term approaches to the TSS include weak selection (relative to local genetic drift; see section 6.5) and weak mutation. Assuming weak selection excludes the ability to study scenarios where selection changes genetic associations and those changes feedback on selection for social traits. These kinds of feedbacks are important both in evolutionary games [48] and in multilocus interactions like the Hill-Robertson effect [74, 95]. The weak mutation assumption of the TSS prevents multiple mutations from competing in a population with the resident phenotype and ensures that the population is fixed for a single phenotype most of the time. Thus, the TSS is inappropriate for studying the evolution of polymorphisms via diversifying or balancing selection. This includes polymorphisms generated by evolutionary branching studied by adaptative dynamics [53, 107]. Adaptive dynamics makes the TSS assumption (at least implicitly) in condition (4) and thus provides at best a necessary condition for branching and polymorphism; typically however, branching is critically dependent on both population size and mutation rate as suggested by the TSS condition [167]. The exclusion of multiple competing mutations under the TSS condition also excludes phenomena such as “clonal interference” [54, 144, 177] that occur when rapid mutations lead to a highly polymorphic population experiencing rapid adaptation [15, 31, 32, 136].

### 6.7. Conclusion

Many of the analytical tools we have used and referenced in this review are much more sophisticated than those readily available to Hamilton when he first discussed inclusive fitness and his eponymous rule [64]. Regardless, the power of the inclusive fitness effect persists both in its ability to suggest broad qualitative patterns connecting natural selection and sociality and in its recurrence as a quantity in complex models of social evolution.

## Acknowledgements

J.V.C. was supported by the National Evolutionary Synthesis Center (NESCent) under NSF grant #EF-0423641.

## Footnotes

1 This condition is very similar to the one obtained in the “strong-selection weak-mutation” limit of Gillespie [p. 221; 56] and the successional-mutations regime of Desai and Fisher [eq. 1; 31]: *N*_T_*μ* log *N*_T_*ω* 1 where *ω* is the strength of selection.

